# Intron 13 retention expands the ADAR1 isoform repertoire to rewire PKR signaling and promote tumorigenesis

**DOI:** 10.64898/2026.05.26.727889

**Authors:** Bryan Y.L. Ng, Jesslyn Zhou, Sze Jing Tang, Yuhua Lim, Vincent Tano, Jovi Jian An Lim, Xi Ren, Omer An, Jinghe Xie, Jian Han, Haoqing Shen, Larry Ng, Wei Liang Gan, Yangyang Song, Ka Wai Leong, Xiaohui Sui, Vanessa Hui En Ng, Jace Koh, Tiffany Ong, Ker-Kan Tan, Leilei Chen

## Abstract

For decades, the biology of ADAR1 has been framed around two major isoforms: the nuclear-enriched p110 and the predominantly cytoplasmic p150. Here, we reveal an unexpected repertoire expansion of ADAR1 isoforms that is driven by intron retention. Specifically, intron 13 (I13) can be retained in *ADAR1* transcripts, conferred by an evolutionarily conserved weaker 5’SS. In addition, we found that the I13 retention (I13R) is negatively autoregulated by ADAR1 through antagonizing the binding of hnRNPA1 to I13 in an editing-independent manner. Despite being sensitive to nonsense-mediated decay, I13R generates two previously uncharacterized truncated isoforms - p90 and p130, that both lack the C-terminal portion of the deaminase domain. Intriguingly, ADAR1p90 – a derivative of the canonical nuclear-enriched p110 isoform – is predominantly cytoplasmic, that effectively represses PKR and eIF2α activation through the sequestration of immunogenic double-stranded RNA (dsRNA) substrates. Furthermore, in a colorectal cancer (CRC) cohort, p90 levels are increased in most tumors relative to matched normal tissues that positively correlates with hnRNPA1 expression. Functionally, xenografts expressing ADAR1p90 grow significantly faster and larger than those expressing ADAR1p110, indicating enhanced tumorigenic potential. These findings revise the canonical view of ADAR1 isoforms, demonstrating that intron retention can generate alternative isoforms with augmented functions that tumors readily exploit.

**One Sentence Summary:** The conserved retention of I13 in ADAR1 transcripts gives rise to previously uncharacterized C-terminal truncated cytoplasmic ADAR1 isoforms, p90 and p130, which, although devoid of catalytic activity, sequester immunogenic dsRNA substrates, thereby preventing PKR binding and downstream activation of eIF2α.

## INTRODUCTION

The adenosine deaminase acting on double-stranded RNA (dsRNA) (ADAR) family comprises ADAR1, ADAR2, and ADAR3, of which only ADAR1 and ADAR2 catalyse all known adenosine-to-inosine (A-to-I) editing events, whereas ADAR3 has no demonstrated catalytic activity (*1–5*). ADAR1 has emerged as a central regulator of dsRNA biology, integrating innate immune and cellular stress pathways triggered by endogenous and exogenous dsRNA. Through A-to-I editing, ADAR1 converts A-U base pairs into less stable I·U wobble pairs, destabilising dsRNA structures that often arise from inverted repeat elements within untranslated regions, including introns and 3’UTRs. In doing so, ADAR1 limits the accumulation of immunostimulatory substrates that would otherwise be sensed by cytoplasmic receptors such as MDA5 and PKR. Beyond its catalytic function, ADAR1 also modulates dsRNA availability through binding and sequestration of dsRNA intermediates, attenuating aberrant activation of PKR signalling(*6–9*).

ADAR1 is expressed as two canonical isoforms, p110 and p150, driven by alternative promoters: a distal constitutive promoter that drives transcription from exon 1B to generate the shorter p110 isoform, and a proximal interferon (IFN)-inducible promoter that drives transcription from exon 1A to produce the full length p150 isoform(*10, 11*). Both isoforms undergo nuclear and cytoplasmic shuttling that depends on nuclear localisation and export signals (NLS and NES)(*12–15*). Notably, an NES within the unique N-terminal Zα domain of p150 is recognised by the export factor exportin 1 (XPO1), facilitating nuclear export and rendering p150 predominantly cytoplasmic, whereas p110 is primarily nuclear. The balance of nuclear import and export can be dynamically modulated by dsRNA abundance and cellular context, further shaping isoform distribution.

Subcellular localisation is a key determinant of ADAR1 function. Nuclear-enriched p110 preferentially edits nascent and structured RNAs, while cytoplasmic p150 positions ADAR1 at the front line of innate sensing to restrain MDA5 and PKR-mediated detection of dsRNA. By limiting MDA5-IRF3 signalling that drives type I IFN responses and preventing PKR-dependent eIF2α phosphorylation that initiates the integrated stress response, ADAR1, particularly p150, acts as a critical brake on cytoplasmic RNA surveillance and maintains immune homeostasis and proteostasis.

In this study, we uncover an unexpected expansion of the ADAR1 isoform repertoire that is driven by intron retention. Specifically, retention of intron 13 (I13R) occurs within *ADAR1* transcripts and is subject to negative autoregulation by ADAR1 itself through a mechanism independent of catalytic editing. Mechanistic investigation reveals that I13R is promoted by the splicing repressor hnRNPA1, which binds to intron 13 of *ADAR1*; however, the canonical ADAR1 isoform p110 counteracts hnRNPA1 binding at its own intron 13, thereby reducing I13R upon ADAR1 overexpression. Although intron 13-retained *ADAR1* transcripts are less stable than canonical isoforms and are sensitive to nonsense mediated decay (NMD), they produce two truncated protein isoforms - p90 and p130, that both lack the C-terminal 120 amino acids (aa) of the deaminase domain. Intriguingly, ADAR1p90 is predominantly cytoplasmic, which suppresses PKR and downstream effector eIF2α phosphorylation by sequestering immunogenic dsRNA substrates. Suppression of the PKR-eIF2α signaling axis lifts translational arrest and blunts pro-apoptotic stress responses, allowing cancer cells to maintain protein synthesis, continue proliferating, and develop larger tumors(*7, 16, 17*). To establish disease relevance, we examined a colorectal cancer (CRC) cohort and found that p90 levels are increased in most CRC tumors relative to matched non-tumor (NT) tissues, with expression potentially correlated with that of hnRNPA1. Consistent with a tumor-promoting role, xenografts expressing p90 grow faster than those expressing p110. Together, these findings revise the prevailing view of ADAR1 isoforms and indicate that intron retention can generate alternative isoforms with functions that tumors readily exploit.

## RESULTS

### ADAR1 negatively regulates intron 13 retention in its own transcripts

Previously, we showed that ADAR1 modulates exon cassette splicing events that influence tumorigenesis(*18*). To identify ADAR1-regulated intron retention (IR) events, we analysed our in-house RNA-seq dataset(*18*) from ADAR1-depleted EC109 cells (esophageal carcinoma cell line) using the IRFinder pipeline(*19–21*). Applying stringent filters (see Materials and Methods), we identified 98 high-confidence IR events reproducible across both biological replicates, of which 84 were downregulated (i.e., reduced IR) and 14 were upregulated upon ADAR1 depletion **(Fig. 1A**). Interestingly, we unexpectedly uncovered the retention of intron 13 (I13R) within *ADAR1* transcripts that is ranked among the top upregulated events under ADAR1-depleted conditions (**Fig. 1A** and **1B**). Indeed, knocking down of ADAR1 in both EC109 and HEK293T cells increased I13R ratio **(Fig. 1C**, **1D** and **S1A**). Moreover, I13R ratio was upregulated in ADAR1 knockout (ADAR1-KO) HEK293T cells compared to wild-type (WT) cells, further corroborating the negative regulation of ADAR1 on its own I13R (**Fig. S1B**). We next constructed an Ex13-I13-Ex14 minigene reporter encompassing exon13-intron13-exon14 of the *ADAR1* gene for further analysis (**Fig. 1E**). As expected, ADAR1 knockdown increased reporter-derived I13R in EC109 cells, and that was rescued upon co-transfection with FLAG-tagged ADAR1p110 **(Fig. 1F** and **1G**). Furthermore, in a separate GFP- fusion reporter whereby intron 13 was inserted into a c-terminal GFP-tagged ADAR1p110 coding sequence, co-expression of FLAG-p110 increased the expression of GFP-p110 fusion protein **(Fig. S1C** and **S1D**). This observation confirms that ADAR1 harbours a repressive role in I13R, which in turn serves as a mechanism of positive feedback regulation on expression of its canonical isoforms. In addition, since no other IR events were observed in *ADAR1* transcripts by our RNA-seq(*18*), we asked why I13R occurs preferentially. Using *in silico* maximum entropy modelling(*22*), we found that the 5′ splice site (5’SS) of intron 13 scores the lowest among all *ADAR1* introns, with deviations from the consensus 5′SS sequence at the +4 and +5 positions (A→C and G→T substitutions, respectively) (**Fig. 1H**). Similarly, the intron 13 3′ splice site (3’SS) ranks among the weakest, indicative of a suboptimal polypyrimidine-tract (**Fig. S1E**). These observations suggest that weak *cis*-spicing motifs in intron 13 likely contribute to its retention within *ADAR1* transcripts. Additionally, we generated a mutant minigene (MUT) by introducing a C>A substitution at +4 position to convert the weak 5’SS into a consensus 5’SS. Indeed, compared to WT minigene, the MUT minigene showed a significant reduction in the I13R ratio (**Fig. 1I**), indicating that cytidine at the +4 position (+4C) likely weakens the intron 13 5’SS. Interestingly, *in silico* comparative genomics analyses using in-built PhyloP and PhastCons pipelines(*23–25*) in the UCSC genome browser revealed that +4C is highly conserved, suggesting that the weakening of intron 13 5′SS and the resulting I13R may have important biological implications (**Fig. 1J**). Separately, qPCR analysis of a commercial multi-tissue cDNA panel revealed I13R across every tissue examined except skeletal muscle, with the spleen harbouring the highest I13R ratio **(Fig. 1K**).

**Figure 1.**
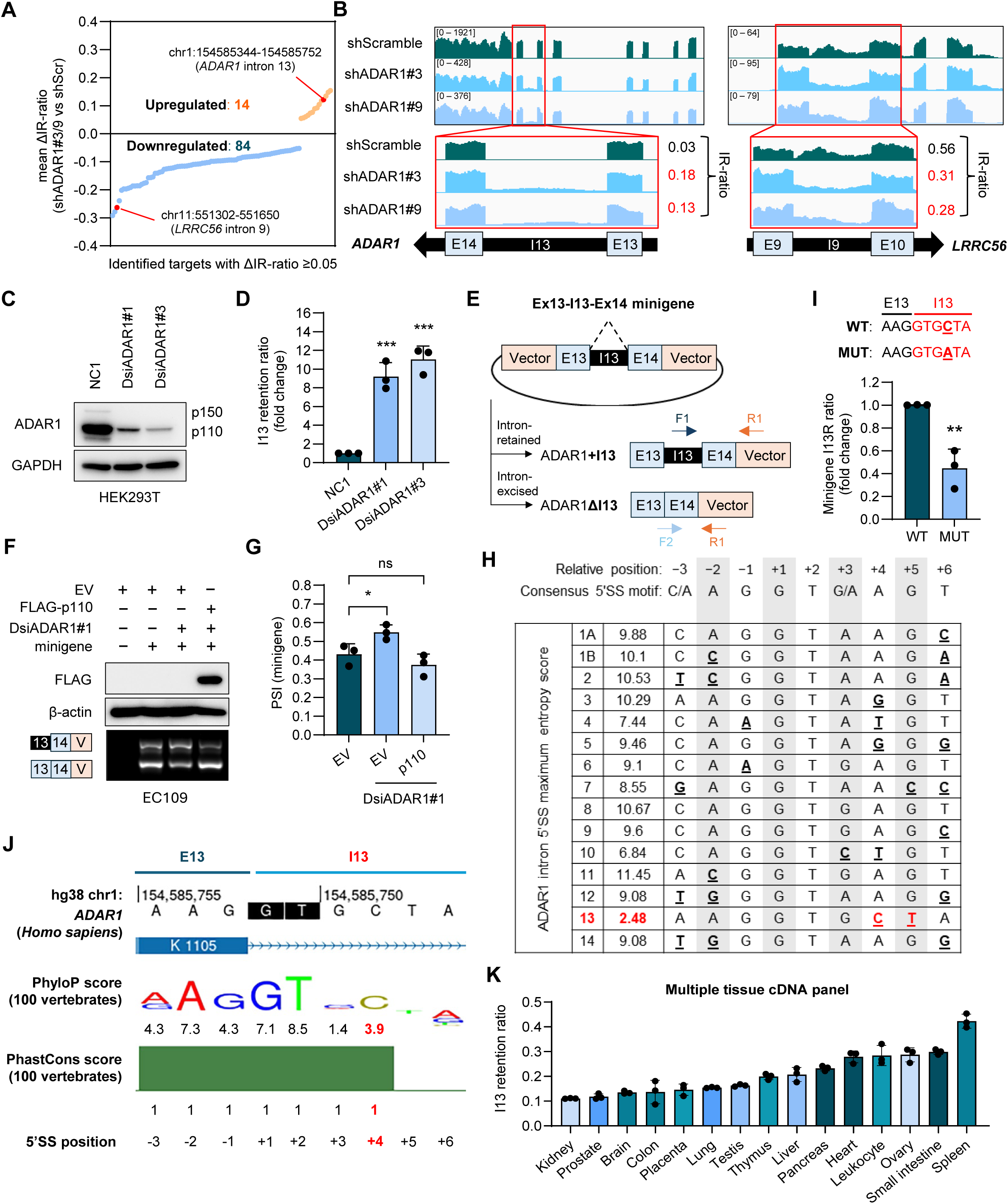
ADAR1 negatively regulates retention of intron 13 in its own transcripts. **(A)** Scatter plot of high-confidence IR events with ΔIR ratio (shADAR1 vs. shScramble) ≥ 0.05 or ≤ −0.05, shared across both shRNAs. **(B)** Integrative Genomics Viewer (IGV) tracks showing RNA-sequencing coverage for the highlighted targets in (**A**). **(C)** Western blot (WB) showing ADAR1 expression in ADAR1 knockdown (KD) and non-targeting control (NC1) HEK293T cells, with GAPDH as the loading control. **(D)** Quantitative PCR (qPCR) quantification of I13R ratios (fold change) in the same groups in (**C**). Fold change is calculated by normalizing the I13R ratio in ADAR1 KD cells to that in NC1 cells. I13R ratios are calculated as described in the Materials and Methods. **(E)** Schematic of the *ADAR1* minigene spanning exon 13-intron 13-exon 14 (Ex13-I13-Ex14), with isoform-specific primers detecting minigene-derived intron 13-retained (ADAR1+I13) and intron 13-excised (ADAR1ΔI13) transcripts: F1, intron 13-specific (dark blue); F2, ΔI13-specific (light blue); R1, common reverse primer targeting the minigene backbone (orange). **(F)** Top, WB analysis of FLAG-tagged ADAR1p110 (FLAG-p110) expression in EC109 cells co-transfected with the minigene described in (**E**) and DsiADAR1 described in (**D**), with β-actin as the loading control. Bottom: RT-PCR detection of minigene-derived *ADAR1+I13* and *ADAR1ΔI13* transcripts in the same groups. **(G)** Quantification of percent spliced-in (PSI) values for minigene-transfected groups in (**F**). **(H)** *In silico* maximum entropy scores for all ADAR1 intronic 5′ splice sites (5’SS). The intron 13 5′SS is highlighted in red, with its deviations from the consensus splice-site motif indicated. **(I)** qPCR quantification of I13R ratios (fold change) of minigene-derived transcripts in HEK293T cells transfected with either the wild-type (WT) Ex13-I13-Ex14 minigene shown in (**E**) or a mutant variant bearing a C>A substitution at the +4 position of the intron 13 5’ splice site (MUT). Fold change is calculated by normalizing the I13R ratio in MUT-transfected cells to that in WT-transfected cells. **(J)** UCSC Genome Browser view of built-in PhyloP and PhastCons conservation scores at the ADAR1 intron 13 5′SS across 100 vertebrate genomes. **(K)** qPCR quantification of I13R ratios in the indicated tissue/cell types. Data are presented as mean ± SD (n = 3 technical replicates). **(D, G, I)** Data are presented as mean ± SD (n = 3 biological replicates). Each data point represents a biological replicate. Statistical significance is determined by two-tailed Student’s *t*-test (unpaired for (**D**) and (**I**); paired for (**G**)) (* *p* < 0.05; ** *p* < 0.01; *** *p* < 0.001; *ns*, not significant).

Together, these findings identify an evolutionarily conserved feature within intron 13 that promotes its retention in *ADAR1* transcripts and show that ADAR1p110 counteracts this retention to enhance its own expression - constituting a positive feedback loop.

### ADAR1p110 represses intron 13 retention by binding to the flanking exons 13 and 14 in an editing-independent manner

Having previously reported that ADAR proteins can regulate cassette exon skipping or inclusion through both editing-independent and -dependent mechanisms(*18*), we asked whether the autoregulatory role of ADAR1 on I13R requires its catalytic (A-to-I editing) activity. To test this, we assessed the effects of a catalytically inactive p110 mutant (DeAD) and a dsRNA-binding null mutant (EAA) on I13R using the Ex13-I13-Ex14 minigene. Notably, the DeAD mutant repressed the I13R ratio to the same extent as WT ADAR1 p110, whereas the EAA mutant significantly attenuated this repression **(Fig. 2A**). These results indicate that regulation of I13R is likely editing-independent but still requires the dsRNA-binding capability of ADAR1.

**Figure 2.**
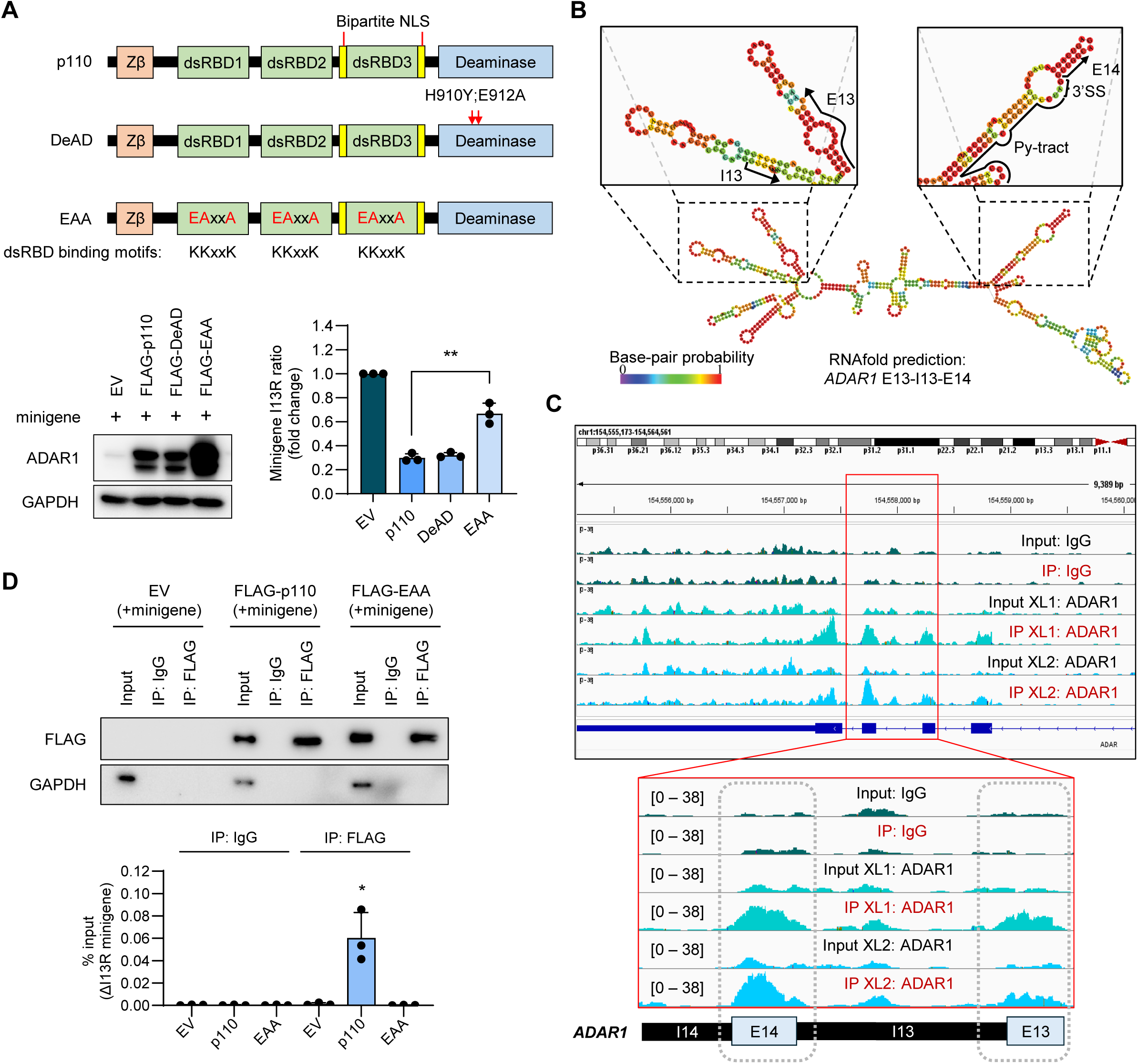
ADAR1p110 represses intron 13 retention by binding to flanking exons 13 and 14 independent of its catalytic ability. **(A)** Top: Schematic of the deaminase-dead (DeAD) and dsRNA-binding–deficient (EAA) mutants with the corresponding mutations indicated (red arrows). Bottom, left: WB analysis of ADAR1 expression in HEK293T cells co-transfected with the Ex13-I13-Ex14 minigene and either WT or mutant ADAR1 constructs; GAPDH serves as a loading control. Bottom, right: qPCR quantification of I13R ratios (fold change) of minigene-derived transcripts in the same groups. Data shown is the mean ± SD (n = 3 biological replicates). Each data point represents one biological replicate. Statistical significance was assessed by two-tailed, unpaired Student’s *t*-test (\*\**p* < 0.01). **(B)** *In silico* prediction of RNA secondary structure within the *ADAR1* exon 13-intron 13-exon 14 region using RNAfold with default parameters. Minimum free energy structure drawings encoding base-pairing probabilities are shown, with base-pairing probabilities indicated by a color spectrum. **(C)** Integrative Genomics Viewer (IGV) tracks at the *ADAR1* locus showing ADAR1 binding peaks from our in-house EC109 ADAR1 eCLIP-seq data. A magnified view of the boxed exon 13-intron 13-exon 14 region is shown below. Data from two technical replicates (XL1 and XL2) are presented. **(D)** RIP-qPCR analysis of the binding of FLAG-tagged ADAR1p110 or the EAA mutant to minigene-derived intron 13-excised *ADAR1* transcripts in HEK293T cells. WB analysis of FLAG-RIP is shown in the top panel. Input corresponds to 5% of total cell lysate. Data are mean ± SD of % input from three biological replicates; statistical significance is determined by a two-tailed, paired Student’s *t*-test (* *p*< 0.05).

Given that the minigene comprises intron 13 flanked by exons 13 and 14, ADAR1 is likely to bind within these regions. RNAfold(*26*) predictions indicated that the exon13-intron13-exon14 segment can form secondary structures despite the lack of repeat elements **(Fig. 2B** and **S2A**). Consistent with this, our in-house ADAR1 eCLIP-seq (enhanced crosslinking and immunoprecipitation followed by RNA sequencing) dataset showed significantly enriched binding peaks across exons 13 and 14 of the *ADAR1* transcript **(Fig. 2C** and **S2B**). Furthermore, RNA immunoprecipitation (RIP) followed by qPCR (RIP-qPCR) confirmed that ADAR1p110 significantly enriched the minigene-derived exogenous ΔI13 (spliced) transcripts, whereas the EAA mutant did not **(Fig. 2D**). Together, these findings suggest that ADAR1p110 binds the flanking exons 13-14 to repress I13R in an editing-independent manner.

### ADAR1p110 counteracts hnRNPA1 association with ADAR1 intron 13 to prevent retention

To further dissect how ADAR1 binding to its own transcript influences I13R, we interrogated our previously published ADAR1 IP-MS (immunoprecipitation followed by mass spectrometry) dataset(*27*) to identify cofactors that may cooperate with ADAR1 to regulate I13R. Relative peptide enrichment in the IP-MS dataset revealed several ADAR1-interacting RNA-binding proteins (RBPs), notably members of the hnRNP and DEAD-box/DHX helicase families, amongst other RBPs with at least 5% enrichment (**Fig. S2C**). Interestingly, publicly available ENCODE eCLIP-seq data(*28*) showed that only hnRNPA1 binds intron 13 in both HepG2 and K562 cells (**Fig. 3A**). We first corroborated the interaction between hnRNPA1 and ADAR1 (**Fig. S2D**). Subsequently, we performed hnRNPA1 RIP, which preferentially enriched minigene-derived intron 13-retained transcripts over the spliced (intron 13-excised, ΔI13) isoform, consistent with intron 13-localized hnRNPA1 peaks observed in ENCODE eCLIP profiles (**Fig. 3A** and **3B**). Functionally, hnRNPA1 knockdown significantly reduced I13R ratio, indicating that hnRNPA1 promotes I13R (**Fig. 3C**). Notably, ADAR1p110 overexpression further suppressed hnRNPA1-induced endogenous I13R, consistent with ADAR1p110 antagonizing the positive regulatory function of hnRNPA1 on I13R (**Fig. S2E**).

**Fig. 3.**
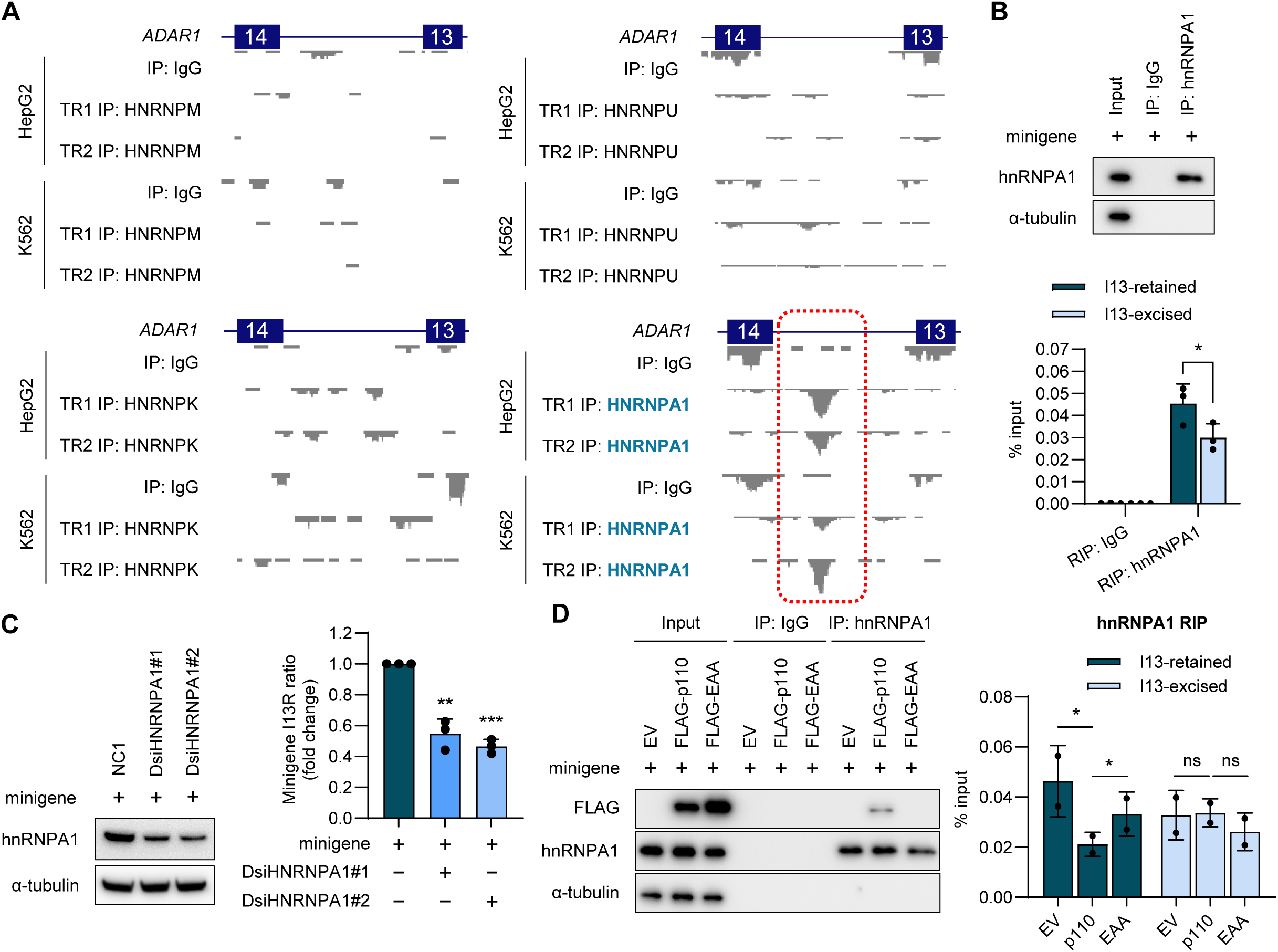
ADAR1p110 antagonizes hnRNPA1 binding to intron 13. **(A)** Genome browser tracks across the *ADAR1* exon 13–exon 14 region showing eCLIP-seq binding peaks for *HNRNPU*, *HNRNPM*, *HNRNPK*, and *HNRNPA1* from publicly available ENCODE datasets. Red box demarcates region of ADAR1 I13 locus where hnRNPA1 binding peaks are called. **(B)** Top: WB analysis of hnRNPA1-RIP immunoprecipitates. Input corresponds to 5% of total cell lysate. Bottom: RIP-qPCR analysis of hnRNPA1 binding to minigene-derived intron 13-retained or - excised *ADAR1* transcripts in HEK293T cells. **(C)** Left: WB analysis of hnRNPA1 expression in HEK293T cells co-transfected with the Ex13-I13-Ex14 minigene and either NC1 or DsiHNRNPA1, with α-tubulin serving as the loading control. Right: qPCR quantification of I13R ratios (fold change) of minigene-derived transcripts in the same groups as (**C**). Data plotted is the mean ± SD (n = 3 biological replicates). Each data point represents one biological replicate. Statistical significance is determined by two-tailed, unpaired Student’s *t*-test (* *p*<0.05; ** *p*< 0.01; *** *p*< 0.001). **(D)** Left: WB analysis of the indicated proteins in hnRNPA1 or IgG immunoprecipitates. Input corresponds to 5% of total cell lysate. Right: RIP-qPCR analysis of hnRNPA1 binding to minigene-derived intron 13-retained or -excised *ADAR1* transcripts in HEK293T cells co-transfected with the Ex13-I13-Ex14 minigene and EV, FLAG-p110, or FLAG-EAA. (**B, D**) Data are presented as mean ± SD of % input (n = 3 biological replicates for (**B**); n = 2 biological replicates for (**D**)). Each data point represents one biological replicate. Statistical significance is determined by a two-tailed, paired ratio *t*-test (* *p*<0.05; ** *p*<0.01; *\*\*\* p*<0.001; *ns*, not significant).

Since ADAR1 binds exons 13-14 to repress I13R (**Fig. 2D**), we hypothesized that ADAR1 may disrupt hnRNPA1-driven I13R by interfering with hnRNPA1 binding to intron 13. To test this, we performed hnRNPA1 RIP in cells transfected with the Ex13-I13-Ex14 minigene (**Fig. 1G**), in the presence of either WT ADAR1 or the EAA mutant overexpression. Strikingly, WT ADAR1 markedly reduced hnRNPA1 enrichment of the intron 13-retained isoform relative to control, whereas the EAA mutant exhibited a much weaker effect, consistent with its impaired binding to dsRNA and hnRNPA1 (**Fig. 3D** and **S2F**). Notably, enrichment of ΔI13 isoform was unchanged, indicating that ADAR1p110 specifically interferes with hnRNPA1 binding to intron 13 and the ensuing I13R (**Fig. 3D** and **S2F**). Altogether, these findings show that ADAR1 binds its own exons 13 and 14 and suppresses I13R in an editing-independent manner by antagonizing hnRNPA1 binding to intron 13.

### Intron 13 retention generates less stable, NMD-susceptible transcripts encoding catalytically inactive ADAR1 p90/p130 isoforms

We next investigated the consequences and fate of I13R in *ADAR1* transcripts. 5′ and 3′ RACE (5′ and 3′ Rapid Amplification of cDNA Ends) assay confirmed the endogenous presence of intron 13-retained *ADAR1p110/p150*-coding transcripts in HEK293T cells (**Fig. 4A**). Since intron 13 spans 408nt and harbors an in-frame premature termination codon (PTC) within its first nine nucleotides, these transcripts are predicted to be substrates of NMD. Indeed, actinomycin D treatment and nascent RNA capture assays showed that intron 13-retained transcripts are less stable than their ΔI13 counterparts (**Fig. S3A** and **S3B**). Consistent with this, UPF1 knockdown significantly increased the abundance of intron 13-retained transcripts together with a known NMD substrate, the exon 4-retained *SRSF4* isoform (**Fig. 4B**), indicating that intron 13-retained transcripts are likely targeted for NMD.

**Fig. 4.**
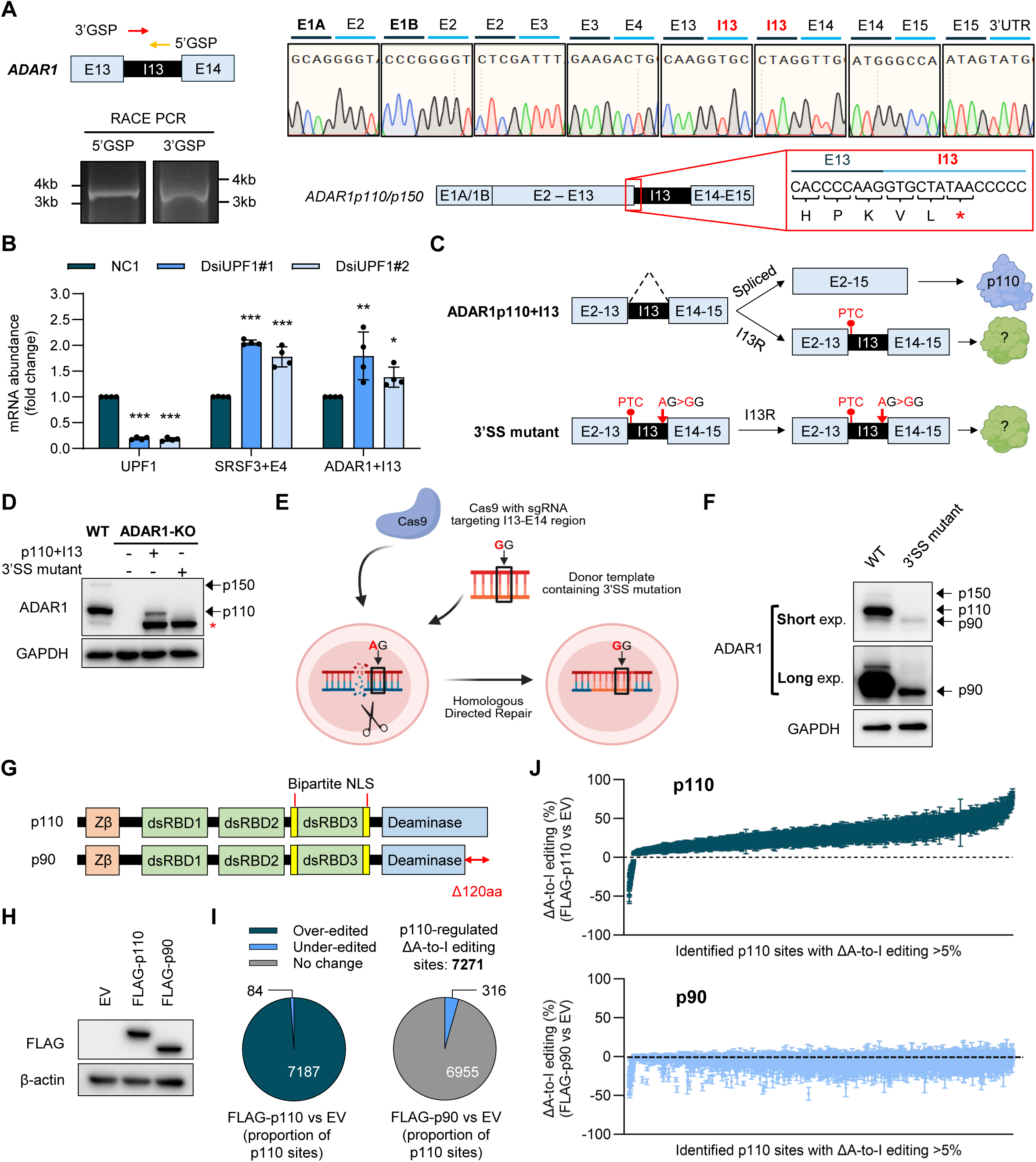
Intron 13 retention generates less stable, NMD-susceptible transcripts encoding catalytically inactive ADAR1 p90/p130 isoforms. **(A)** Top, left: schematic of 5′ and 3′ gene-specific primers (5′GSP and 3′GSP, respectively) targeting *ADAR1* intron 13. Bottom, left: PCR amplicons generated with the indicated GSPs, resolved by agarose gel electrophoresis. Top, right: Sanger sequencing chromatograms showing exon-exon or exon-intron junctions of cloned RACE-PCR products. Bottom right: schematic illustrating intron 13 retention in p110- or p150-encoding transcripts. The exon 13-intron 13 boundary is highlighted with a red box, showing the translated amino acid sequence and a premature termination codon (PTC) marked with a red asterisk. **(B)** RT-qPCR analysis of mRNA levels (fold change) for the indicated transcripts in HEK293T cells upon UPF1 KD. Fold change is calculated by normalizing mRNA levels in UPF1 KD cells to NC1 controls. Data are presented as mean ± SD from four biological replicates. Statistical significance is determined by two-tailed, unpaired Student’s *t*-test (* *p*< 0.05; ** *p*< 0.01; *** *p*< 0.001). **(C)** Schematic of the ADAR1p110+I13 construct, in which intron 13 is inserted between exons 13 and 14 of an ADAR1p110 expression plasmid. Splicing of the intron yields canonical p110, whereas intron retention produces a truncated, noncanonical ADAR1 isoform. The 3′SS mutant is identical to ADAR1p110+I13 except for an AG→GG substitution at the 3′SS to ablate splicing, thereby enforcing intron retention and translation of the truncated isoform. **(D)** WB analysis of ADAR1 isoforms in WT HEK293T cells and ADAR1-KO HEK293T cells transduced with the constructs described in (**C**). The asterisk denotes the truncated ADAR1 isoform. **(E)** Schematic of CRISPR-Cas9-mediated homology-directed repair (HDR) to ablate the *ADAR1* intron 13 3′SS in HEK293T cells. **(F)** WB analysis of endogenous p90 in 3′SS mutant cells generated as described in (**E**). **(G)** Schematic illustrating the C-terminal 120aa truncation within the deaminase domain of ADAR1p90. **(H)** WB analysis of the FLAG-tagged protein expression in HEK293T cells overexpressed with FLAG-p110 and FLAG-p90, with β-actin as the loading control. **(I)** Left: Pie chart of the 7,271 high-confidence p110-regulated A-to-I editing sites identified by RNA-seq. Sites are classified as over-edited, under-edited, or no change relative to control. Right: Pie chart showing how these same 7,271 sites behave upon p90 overexpression, categorized as over-edited, under-edited, or no change using the same filter criteria. **(J)** Waterfall scatter plot showing changes in A-to-I editing efficiency (ΔA-to-I editing, %) at 7,271 ADAR1p110-regulated sites, rank-ordered by effect size, in cells overexpressing FLAG-ADAR1p110 (top) or FLAG-ADAR1p90 (bottom) relative to EV control. Only sites exhibiting ≥5% change in editing level upon overexpression and supported by a minimum read coverage of 20 reads were considered as regulated sites and included in the analysis. Error bars represent variability across replicates.

We next asked whether these transcripts can be translated despite NMD. To test this, we engineered an ADAR1p110+I13 construct by inserting intron 13 between exons 13 and 14 of an ADAR1p110 expression plasmid and introduced an AG→GG mutation at 3′ SS (3′SS mutant) to prevent intron 13 excision (**Fig. 4C**). Intriguingly, in ADAR1-KO HEK293T cells expressing the ADAR1p110+I13 construct, we detected both canonical p110 which arises from intron 13 removal and a truncated ∼90 kDa ADAR1 isoform (**Fig. 4D** and **S3C**). In contrast, cells expressing the 3′SS mutant produced only the truncated isoform (**Fig. 4D** and **S3C**). To corroborate these findings in an endogenous context, we used CRISPR-mediated homology-directed repair (HDR) to generate cells harboring the 3′SS mutation (**Fig. 4E**). Consistently, a truncated ∼90 kDa ADAR1 isoform (hereafter referred to as p90) was readily detected in the 3′SS mutant cells and was modestly detectable in WT cells (**Fig. 4F**). Moreover, ADAR1 IP from 3′SS mutant cells enriched an additional ∼130 kDa isoform, likely derived from the intron 13-retained p150 transcript (hereafter p130) (**Fig. S3D**). Given that hnRNPA1 promotes I13R **(Fig. 3**), we next tested whether altering hnRNPA1 levels modulates p90 expression. Indeed, hnRNPA1 knockdown reduced ADAR1p90 in 3′SS mutant cells (**Fig. S3E**).

Since ADAR1p90 is truncated by ∼120aa at the C-terminus within the deaminase domain (**Fig. 4G**), we next asked whether p90 retains catalytic activity. Overexpression of p110 or p90 in HEK293T cells followed by deep RNA sequencing (RNA-seq) identified 7,271 and 937 high-confidence A-to-I editing sites regulated by p110 and p90, respectively, with each site showing a mean editing change of ≥5% versus control **(Fig. 4H**, **4I** and **Fig. 3F**). Among the 7,271 p110-regulated sites, the vast majority (7,187; 98.8%) were over-edited **(Fig. 4H** and **4I**) (hereafter referred to as ‘p110 over-edited sites’). Strikingly, none of these sites showed a ≥5% increase in editing frequency upon p90 overexpression; instead, 95.7% (6,955) exhibited no change and 4.3% (316) decreased in editing, suggesting a potential dominant-negative effect on endogenous ADAR1 (**Fig. 4J**). Further analysis of the 937 p90-regulated sites revealed that all sites exhibited decreased editing frequencies upon p90 overexpression, whereas only a small subset (43; 4.6%) showed a corresponding decrease with p110 overexpression (**Fig. S3F**), confirming the dominant-negative effect of p90 on endogenous ADAR1 editing activity. One plausible mechanism is that p90, lacking full catalytic capacity, competes for dsRNA substrates and/or dimerization interfaces, thereby impairing editing mediated by canonical ADAR1. Collectively, these findings indicate that although *ADAR1p90/p130* transcripts are relatively more unstable than canonical *p110/p150* transcripts owing to NMD, the limited efficiency of this pathway allows translation of truncated, catalytically inactive isoforms. These isoforms may exert dominant-negative effects by outcompeting canonical ADAR1 for dsRNA binding and/or by disrupting ADAR1 homodimer formation, thereby suppressing RNA editing.

### ADAR1p90 represses PKR activation through cytoplasmic dsRNA sequestration

Recent advances have established ADAR1 as a suppressor of dsRNA-directed innate immunity (*6, 17, 29, 30*). Through A-to-I RNA editing, ADAR1 marks endogenous RNAs as “self,” preventing their aberrant recognition by the cytoplasmic sensors MDA5 and PKR. More recently, the IFN-induced ADAR1p150 isoform was shown to repress PKR activation in an editing-independent manner by sequestering dsRNA intermediates, thereby preventing PKR engagement (*6–9*).

Given that the catalytically inactive ADAR1p90 retains all functional domains except the C-terminal 120aa of the deaminase domain, we investigated its role in dsRNA sensing by MDA5/PKR and their downstream signaling. Since HEK293T cells exhibit low basal expression of MDA5 and PKR (*9, 30*), we treated cells with IFNβ to elevate their levels. IFNβ increased total and phosphorylated PKR as well as MDA5, but only modestly activated downstream effectors, including eIF2α via PKR and IRF3 via MDA5 (lane 2 vs. lane 1, **Fig. 5A** and **5B**). Transient ADAR1 knockdown exacerbated PKR-eIF2α and MDA5-IRF3 activation, as evidenced by markedly higher phospho-to-total ratios for eIF2α and IRF3 (lane 5 vs. lane 2, **Fig. 5A** and **5B**). Strikingly, overexpression of either ADAR1p110 or ADAR1p90 repressed PKR activation, with ADAR1p90 producing significantly stronger suppression of PKR and eIF2α phosphorylation than p110 despite its catalytic inactivity (lanes 4 and 3 vs. 2; lanes 7 and 6 vs. 5, **Fig. 5A** and **5B**). In contrast, neither isoform showed consistent repression of MDA5-IRF3 activation (lanes 4 and 3 vs. 2; lanes 7 and 6 vs. 5, **Fig. 5A** and **5B**).

**Fig. 5.**
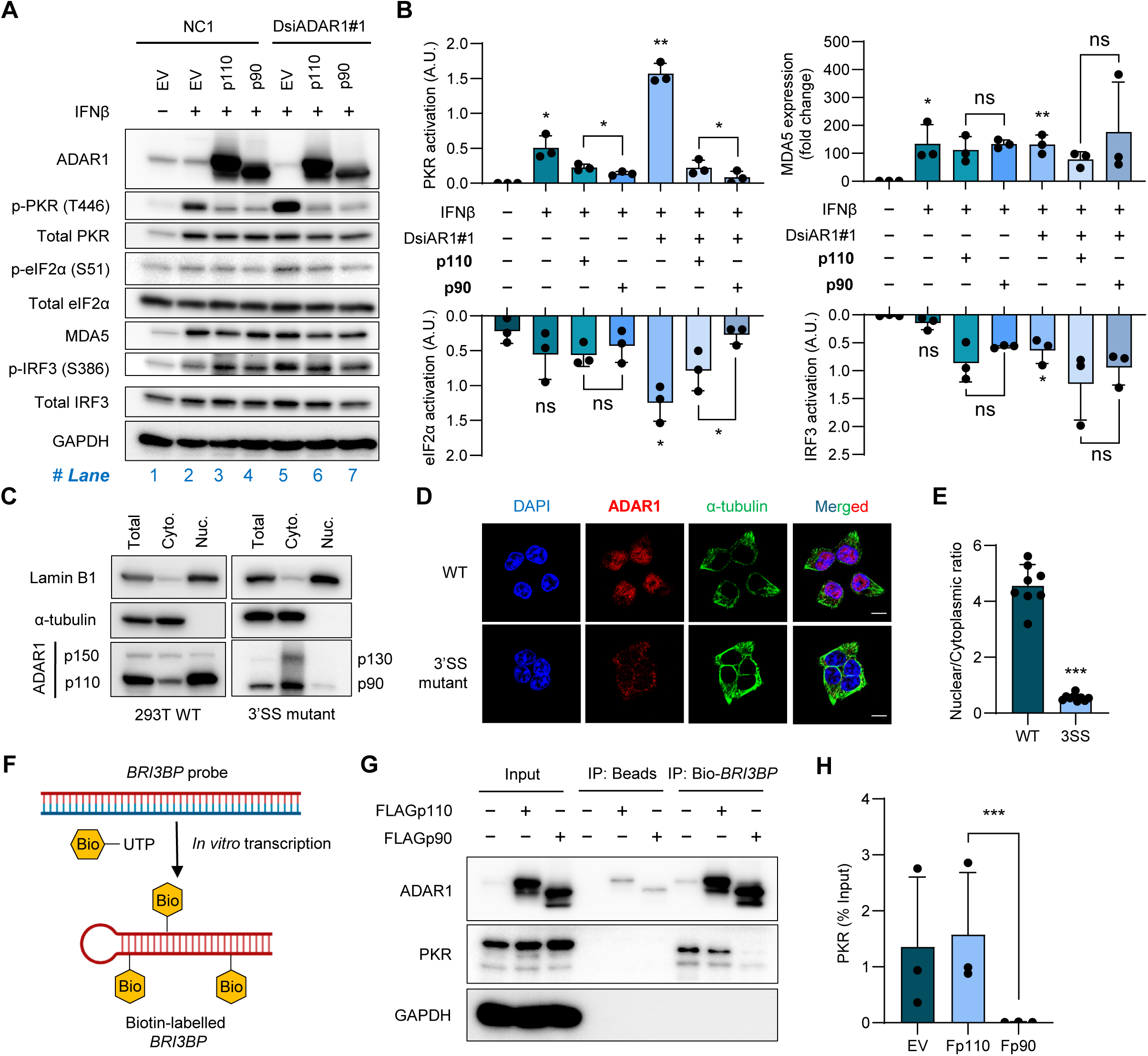
ADAR1p90 represses PKR-eIF2α activation through cytoplasmic dsRNA sequestration. **(A)** WB analysis of the indicated proteins in HEK293T cells stimulated with IFN-β and subjected to transient ADAR1 KD (DsiADAR1#1), followed by rescue with EV, p110, or p90. The first lane (NC1, EV, no IFN-β) serves as an unstimulated baseline control. GAPDH serves as the loading control. **(B)** Quantification of p-PKR/total PKR, p-eIF2α/total eIF2α, and p-IRF3/total IRF3 band intensity ratios (reflecting PKR, eIF2α, and IRF3 activation, respectively), as well as MDA5 band intensity normalized to GAPDH (expressed as fold change), measured using ImageJ. Phosphorylation ratios are expressed in arbitrary units (A.U.). Data are presented as mean ± SD from three independent biological replicates (n = 3). Statistical significance was determined by two-tailed paired Student’s *t*-test, except for MDA5 expression (fold change), which was assessed by a two-tailed unpaired Student’s *t*-test (* *p*<0.05; ** *p*<0.01; *ns*, not significant). A representative western blot is shown in (**A**). **(C)** WB analysis of Lamin B1 (nuclear marker), α-tubulin (cytoplasmic marker), and ADAR1 isoforms in total lysates and nuclear (Nuc.) and cytoplasmic (Cyto.) fractions from WT (expressing p150 and p110) and 3′SS mutant (expressing p130 and p90) HEK293T cells. **(D)** Immunofluorescence (IF) staining of ADAR1 (red) and α-tubulin (green) in WT and 3′SS mutant HEK293T cells. Nuclei are counterstained with DAPI (blue). The merged panel shows all three channels. Images are representative of multiple fields of view (FOVs) from three independent experiments. Scale bar, 10 μm. **(E)** Quantification of the nuclear-to-cytoplasmic ratio of ADAR1 fluorescence intensity in WT and 3′SS mutant HEK293T cells. Data are presented as mean ± SD from 8–9 FOVs per condition. Statistical significance was determined by a two-tailed unpaired Student’s *t*-test (*** *p*< 0.001). **(F)** Schematic of the biotin-labelled *BRI3BP* dsRNA probe generated by *in vitro* transcription incorporating biotin-UTP. The probe corresponds to the long dsRNA structure within the 3′ UTR of *BRI3BP*. **(G)** WB analysis of ADAR1, PKR, and GAPDH in pulldown products captured by the biotin-labelled *BRI3BP* probe from HEK293T cells expressing FLAG-p110 or FLAG-p90. A beads-only pulldown (no probe) serves as the negative control. GAPDH, detected in input but absent in pulldown fractions, confirms the specificity of probe-dependent enrichment. Input corresponds to 5% of total cell lysate. **(H)** Quantification of PKR band intensities from (**G**), measured using ImageJ and normalised to input. Data are presented as mean ± SD from three independent biological replicates (n = 3). Statistical significance was determined by a two-tailed paired ratio *t*-test (*** *p* < 0.001).

To explain why ADAR1p90 represses PKR activation more effectively than canonical ADAR1p110, we propose that p90 competes with PKR for binding to immunogenic dsRNA substrates, analogous to the mechanism reported for ADAR1p150(*8*). Although truncation of the C-terminal 120 amino acids was not predicted to disrupt the bipartite nuclear localization signal (NLS) flanking the third dsRNA-binding domain (dsRBD3)(*31*), cellular fractionation of HEK293T WT and 3′SS mutant cells revealed that ADAR1p90 is predominantly cytoplasmic, in contrast to the nuclear-enriched ADAR1p110 (**Fig. 5C**). Consistent results were observed in HEK293T cells upon overexpression of MYC-tagged ADAR1p90 or p110 (**Fig. S4A**). Immunofluorescence (IF) analysis corroborated these findings (**Fig. 5D** and **5E**), suggesting that, contrary to prior assumptions, a putative NLS (or nuclear retention element) may reside within the deleted C-terminal portion of the deaminase domain. This cytoplasmic enrichment of p90 provides a plausible basis for its enhanced repression of PKR activation via dsRNA sequestration.

In addition to its cytoplasmic enrichment, we examined whether ADAR1p90 displays higher binding affinity for cytoplasmic immunogenic dsRNA than ADAR1p110. To identify a suitable immunogenic dsRNA target, we integrated three datasets: (1) our ADAR1p150 eCLIP-seq dataset to detect cytoplasmic ADAR1-bound dsRNA loci, (2) a published MDA5 protection assay mapping transcriptome-wide MDA5 binding sites to identify potential immunogenic dsRNA substrates(*32*), and (3) our in-house cytoplasmic J2 (anti-dsRNA) eCLIP-seq dataset to profile dsRNA abundance (**Fig. S4B**). From this analysis, we selected *BRI3BP1* as an exemplary target harboring a long dsRNA structure within its 3′ UTR (**Fig. S4C**). A biotin-labelled dsRNA probe corresponding to this *BRI3BP1* element was generated by *in vitro* transcription incorporating biotin-UTP (**Fig. 5F**). In biotin-RNA pulldown assays, expression of ADAR1p90, but not ADAR1p110, markedly reduced PKR binding to the biotin-labelled *BRI3BP1* probe, suggesting that ADAR1p90 competes with PKR for occupancy at this immunogenic dsRNA substrate (**Figs. 5G** and **5H**).

Together, these results show that ADAR1p90 suppresses PKR-eIF2α signalling activation more potently than ADAR1p110 via an editing-independent mechanism, acting as a cytoplasmic dsRNA decoy that sequesters immunogenic substrates away from PKR.

### ADAR1p90 is upregulated in colorectal cancer and promotes tumorigenicity

Repression of PKR-eIF2α signaling relieves translational arrest and dampens pro-apoptotic stress responses, enabling cancer cells to sustain protein synthesis, proliferate, and form larger tumors(*7, 16, 17*). Given that ADAR1p90 harbors an augmented capacity to repress PKR-eIF2α signalling, we investigated whether intron 13-retained *ADAR1* transcripts are upregulated in cancer. In a cohort of 32 matched colorectal cancer (CRC) and adjacent non-tumor (NT) tissue samples, CRC tumors exhibited significantly higher I13R ratios accompanied by elevated total *ADAR1* expression compared with their NT counterparts (**Fig. 6A**). To further assess ADAR1p90 protein levels, we selected six representative matched pairs showing the largest differences in I13R ratio between CRC and their respective NT samples. Notably, in four cases (P35, P43, P48, and P60) with markedly elevated hnRNPA1 expression, CRC tumors displayed higher ADAR1p90-to-p110 ratios, whereas the remaining two cases (P40 and P66), in which hnRNPA1 expression was decreased or only modestly increased, showed correspondingly low ADAR1p90 levels (**Fig. 6B**). Together, these findings validate the positive regulatory role of hnRNPA1 in promoting intron 13 retention in clinical CRC specimens and highlight the potential relevance of this splicing event in cancer progression.

**Fig. 6.**
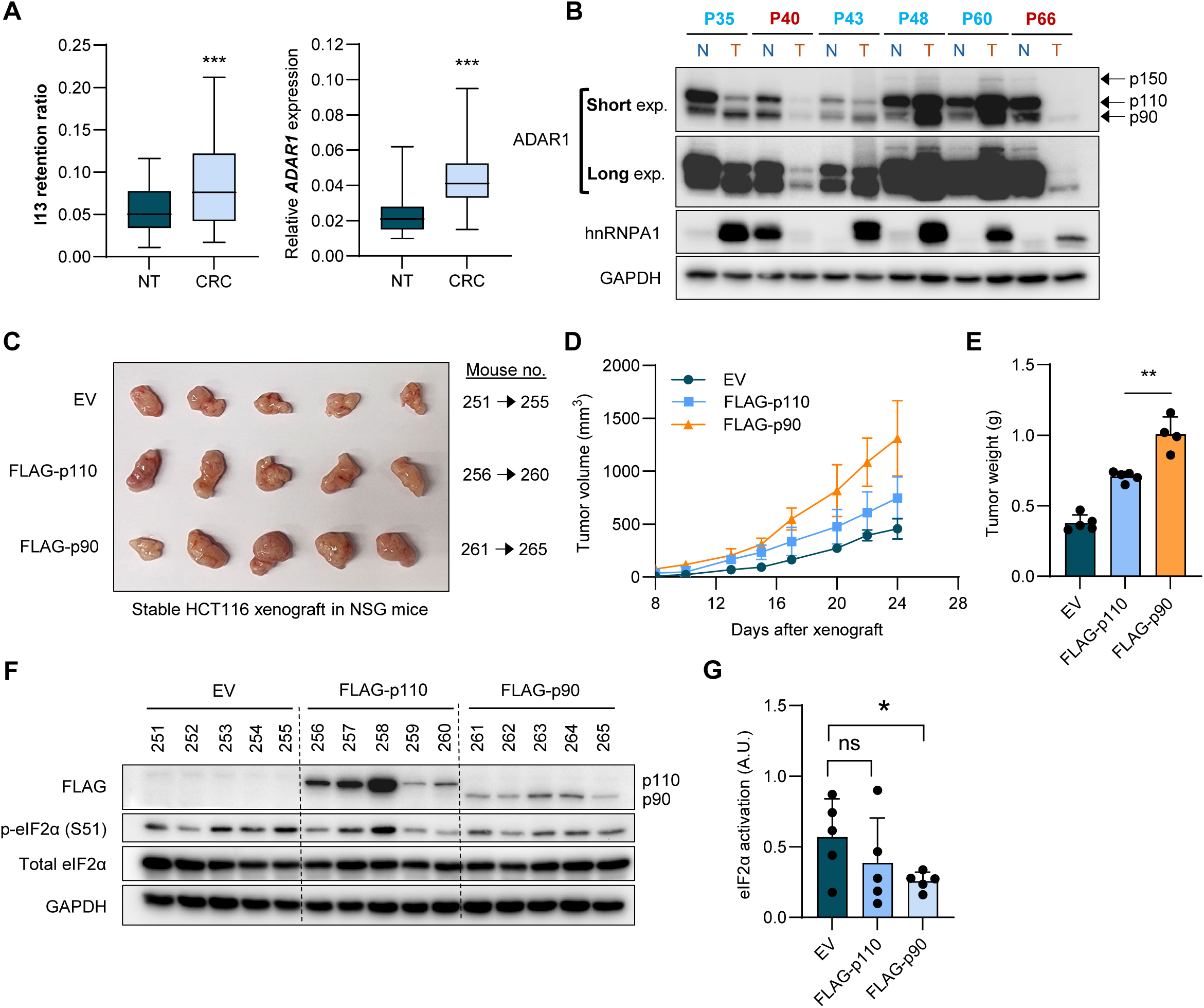
ADAR1p90 is upregulated in colorectal cancer and promotes tumorigenicity. **(A)** RT-qPCR analysis of the I13R ratio and total *ADAR1* transcript levels (relative expression [ΔΔC_t_]) in 32 matched pairs of CRC tumor and adjacent non-tumour (NT) tissue samples. Data are presented as mean ± SD (*n* = 32 pairs). **(B)** WB analysis of ADAR1 isoforms (p130 and p90), hnRNPA1, and GAPDH in six representative matched CRC tumor-NT pairs selected on the basis of the largest differences in I13R ratio among the 32 pairs. **(C)** Representative images of subcutaneous xenograft tumours derived from HEK293T cells stably expressing the indicated constructs, harvested at experimental endpoint. **(D)** Xenograft tumor growth curves over a 24-day observation period. Data are presented as mean ± SD (*n* = 5 tumors per group), except for the FLAG-ADAR1p90 group (*n* = 4), from which one tumor (mouse 261) was excluded. **(E)** Endpoint tumor mass quantified by digital weighing scale. Each data point represents an individual tumor. **(F)** WB analysis of p-eIF2α, total eIF2α, and the indicated proteins in lysates prepared from endpoint xenograft tumors. **(G)** Quantification of the p-eIF2α-to-total eIF2α band intensity ratio from the western blots shown in (**F**), as measured by densitometry using ImageJ. Arbitrary units (A.U.) were calculated as described in **Fig. 5B**. **(A, E, G)** Statistical significance is determined by a two-tailed, paired Student’s *t*-test (* *p*< 0.05; ** *p*< 0.01; *** *p*< 0.001; *ns*, not significant).

We next asked whether ectopic overexpression of ADAR1p90 in the CRC cell line HCT116 promotes xenograft growth to a greater extent than its p110 counterpart. In four of five mice per group, p90-expressing tumors grew significantly faster than those in the control or p110 groups, and endpoint tumor weights were significantly higher for p90-expressing tumors than for EV control- or p110-derived tumors **(Fig. 6C-E**). Given that ADAR1p90 repressed PKR-eIF2α activation **(Fig. 5**), we evaluated this pathway in endpoint xenografts. Expectedly, eIF2α activation was significantly repressed compared with the control group, potentially accounting for the heightened tumorigenicity of p90-expressing tumors **(Fig. 6F** and **6G**).

Collectively, these findings establish disease relevance by showing that p90-to-p110 ADAR1 expression ratios are elevated in CRC tumors compared with matched non-tumor tissues and positively correlate with hnRNPA1 levels. Furthermore, this indicates that ADAR1p90 possesses greater oncogenic potential than ADAR1p110, possibly through repression of PKR-eIF2α signaling.

## DISCUSSION

Although ADAR2 has been shown to undergo autoregulated alternative splicing where ADAR2-mediated editing within its own pre-mRNA creates a proximal 3’ splice site that inserts 47 nucleotides into the coding region, producing a frameshift and a truncated, inactive protein(*33*), there have, to date, been no reports of an autoregulatory mechanism for ADAR1 via alternative splicing. For decades, the biology of ADAR1 has been centered on two major isoforms: the nuclear-enriched p110 and the predominantly cytoplasmic p150. Here, we identify a previously unreported autoregulatory mechanism in which ADAR1 p110 suppresses I13R in its own transcripts. Our findings reveal an evolutionarily conserved *cis*-regulatory feature within intron 13 that promotes selective intron retention. Intron 13 harbors a weakened 5′SS marked by a non-consensus cytidine at the +4 position, which is highly conserved across vertebrates, instead of the canonical adenosine. Correcting this nucleotide to match the consensus 5′SS markedly improved intron 13 removal, indicating that the +4 position is critical for exon–intron definition. Notably, cytidine at +4 does not base-pair with U1 snRNA (*34–36*), and this diminished interaction likely weakens 5′SS recognition, thereby favoring I13R. Besides, we identify hnRNPA1 as a positive regulator of I13R, however, ADAR1 binds its own exons 13 and 14 to interfere with hnRNPA1 association at intron 13.

Several questions remain: How prevalent is ADAR1 I13R across tissues and cell types, and under which physiological or pathological conditions are intron 13-retained transcripts increased? Here, we show that while I13R is detectable across multiple tissues and cell types - leukocytes, ovary, small intestine and most notably the spleen, exhibit higher I13R ratios than other tissues. This is consistent with a landmark study demonstrating that increased IR-NMD coupling in several functional genes is essential for shaping nuclear architecture as myeloid progenitors commit to granulopoiesis(*21*). Accordingly, the elevated I13R ratio in spleen may reflect its immune-cell-dense environment and substantial granulocyte population. Notably, the same study identified *ADAR1* among 855 genes with at least one retained intron in human granulocytes, although the specific intron was not specified(*21*). Whether granulocytes indeed express higher levels of ADAR1 p90 or p130 than canonical isoforms to support their function and survival remains to be determined.

As a positive regulator of ADAR1 I13R, hnRNPA1 has been implicated in cellular differentiation across diverse lineages, including granulocytes(*37*) and cardiomyocytes(*38*), through its roles in gene expression and alternative splicing. Canonical ADAR1 expression declines during terminal differentiation(*39*), but whether this reduction reflects epigenetic silencing, I13R-NMD coupling, or additional regulatory layers is unresolved. In contrast, ADAR1 is frequently upregulated in cancer(*40–53*), consistent with the de-differentiated state that is characteristic of many tumors. It is therefore plausible that initial ADAR1 elevation during early de-differentiation is further reinforced by its positive autoregulatory suppression of I13R. However, in cellular contexts with high hnRNPA1 expression, such as our CRC cohort, this circuit may be attenuated: because hnRNPA1 promotes I13R, it can shift the balance toward production of ADAR1 p90 rather than canonical ADAR1, and ADAR1 p90 displays tumorigenic potential in our xenograft models. Although intron 13-retained transcripts are subject to NMD, a fraction escapes degradation and is translated into truncated, catalytically inactive ADAR1 isoforms. A recent single-cell study showed that cells within a population vary in SMG1 abundance - the kinase that phosphorylates UPF1 to activate NMD(*54*). Cells with lower SMG1 likely exhibit attenuated NMD, enabling PTC-containing transcripts to escape decay, providing a plausible basis for detectable ADAR1 p90 expression in subpopulations with reduced SMG1. Independently, extensive work demonstrates that NMD is attenuated during physiological stress, allowing translation of otherwise NMD-sensitive targets(*55*). Under oxidative stress, heat shock, viral infection, or inflammatory exposure, cells experience endoplasmic reticulum (ER) stress and activate the unfolded protein response (UPR). A central consequence of the UPR is the phosphorylation of eIF2α, which suppresses global translation to alleviate ER burden. Because NMD is translation-dependent, it pauses under these conditions, permitting selective translation of stress mediators such as ATF3, ATF4, and CHOP(*56–58*). Within this framework, ER stress or dsRNA-PKR sensing would activate eIF2α, inhibit NMD, and consequently increase ADAR1 p90 expression. ADAR1 p90 could then exert a pro-survival role by repressing PKR-eIF2α signaling to restore protein synthesis and support cellular homeostasis. In tumors, where nutrient scarcity, hypoxia, and reactive oxygen species are pervasive, eIF2α-mediated NMD shutdown is commonly observed(*59–65*). Thus, the elevated ADAR1 p90-to-p110 ratios in our CRC cohort may reflect NMD attenuation coupled with increased hnRNPA1 expression.

In summary, we uncover a previously unreported negative autoregulatory loop in which ADAR1 p110 suppresses intron 13 retention by antagonizing hnRNPA1 binding to intron 13, and we provide an example of intron retention generating alternative isoforms with augmented functions relative to canonical counterparts. We further show that a conserved, weakened 5′ splice site positions intron 13 for retention; under stress-induced NMD shutdown, this facilitates translation of ADAR1 p90 as part of the integrated stress response - a pro-survival mechanism that colorectal tumor cells may exploit under chronic stress conditions.

## ACKNOWLEDGEMENTS

We thank and acknowledge the Microscopy and Multiplex Assays (MMA) Core Facility at the Cancer Science Institute of Singapore for their assistance with imaging experiments. This work was supported by National Research Foundation Singapore; Singapore Ministry of Education under its Research Centres of Excellence initiative; Singapore Ministry of Education’s Tier 2 Grant [MOE- T2EP30123-0003] and Tier 3 Grant [MOET32023-0003]; NMRC Clinician Scientist-Individual Research Grant (project ID: MOH-001092-00); NMRC Open Fund-Large Collaborative Grants (project IDs: MOH-001573-00 and MOH-001067); and NRF Competitive Research Programme (CRP) Grants (Grant numbers: NRF-CRP26-2021-0001 and NRF-CRP26-2021-0005).

## AUTHOR CONTRIBUTIONS

B.Y.L.N. and L.C. conceptualized the study. B.Y.L.N. performed most of the experiments. S.J.T., J.Z. and Y.L. performed experiments that are part of this study. V.T., J.J.A.L. and B.Y.L.N. conducted the bioinformatic analyses. O.A., R.X. and J.X. executed the IRFinder pipeline analyses. B.Y.L.N., Y.S., K.W.L., X.S. and J.Z. processed and performed experiments related to CRC clinical samples. V.H.E.N. provided materials and assisted with mice experiments. J.H., H.S., L.N., W.L.G., Y.S., J.K. and T.O. provided insightful suggestions. B.N. and L.C. wrote the manuscript. L.C. supervised the study.

## MATERIALS AND METHODS

### Cell culture

HEK293T (ATCC CRL-3216) and HCT116 (ATCC CCL-247) cell lines were obtained from the American Type Culture Collection (ATCC). EC109 cells were kindly provided by Professor George Sai Wah Tsao (Director, Faculty Core Facility, Li Ka Shing Faculty of Medicine). A CRISPR-mediated ADAR1 knockout HEK293T cell line was generated in-house. HEK293T 3’SS mutant cells harboring an A>G mutation at the GRCh38.p14 chr1:154585346 locus were generated by Creative Biogene. All cells were maintained at 37°C in a humidified incubator with 5% CO₂. HEK293T cells were cultured in Dulbecco’s Modified Eagle Medium (DMEM; Gibco, 11965092) supplemented with 10% fetal bovine serum (FBS; Gibco, 10438026). EC109 cells were cultured in RPMI 1640 medium (Gibco, 11875093) supplemented with 10% FBS. HCT116 cells were cultured in McCoy’s 5A Modified Medium (Gibco, 16600082).

### Plasmid construction

Minigene sequences were amplified from human placenta genomic DNA (Sigma) using PrimeSTAR Max DNA polymerase (Takara) and subcloned into pcDNA3.1+ plasmid. Coding sequences of proteins were amplified with Platinum SuperFi II PCR Polymerase (Invitrogen, 12369010) and cloned into Lenti-X Expression System (EF1alpha Version) (Takara, 631253), pEGFP-N1 or pcDNA3.1+ (with c-terminal MYC tag). Insertion of intron 13 into ADAR1p110-coding plasmid was performed using the overlap extension PCR and recombination protocol(*66*). Site-directed mutagenesis was carried out using primers containing respective mutations according to the protocol of the manufacturer of the Platinum SuperFi II PCR polymerase.

### Transfection

HEK293T or EC109 cells were seeded in 6-well plates and allowed to incubate at 37°C in a humidified incubator with 5% CO_2_ for 24 hours prior to transfection. In brief, for minigene assays, cells were transfected with 500ng minigene for no more than 24 hours using lipofectamine 2000 (Invitrogen, 11668027) according to the manufacturer protocol. In co-expression minigene models, cells were transfected with 25pmol pre-designed dicer-substrate siRNA (DsiRNA) from Integrated DNA Technologies (IDT) [Design IDs: DS NC1, hs.Ri.ADAR.13.1/3 (#1/#3), hs.Ri.HNRNPA1.13.1/2 (#1/#2), hs.Ri.UPF1.13.1/2 (#1/#2)] using RNAiMAX reagent (Invitrogen, 13778075) or 1ug expression plasmid for 24 hours, followed by a subsequent transfection of minigene for another 24 hours incubation prior to harvesting for minigene assays. For rescue minigene models, DsiRNA and rescue expression plasmid was complexed separately in RNAiMAX or Lipofectamine 2000, respectively, according to the manufacturer protocol prior to transfection.

### CRISPR-mediated introduction of the 3′ splice site mutation

To introduce the 3′ splice site (3′SS) mutation at the intron 13-exon 14 (I13-E14) boundary of the endogenous *ADAR1* locus, CRISPR-Cas9-mediated homology-directed repair (HDR) was employed. A single guide RNA (sgRNA) targeting the I13-E14 region was designed and cloned into an appropriate Cas9 expression vector. The sgRNA-Cas9 ribonucleoprotein complex was co-transfected into HEK293T cells together with a donor template harboring the desired 3′SS point mutation, in which the endogenous splice acceptor sequence (AG) was substituted with a non-canonical dinucleotide (GG) to disrupt canonical splicing at this site. Following transfection, cells were allowed to recover and undergo HDR-mediated incorporation of the donor template. Successfully edited clones were selected and screened by PCR amplification and Sanger sequencing across the I13-E14 junction to confirm precise introduction of the 3′SS mutation. Homozygous or heterozygous mutant clones were expanded for downstream functional analyses.

### IRFinder analyses

The EC109 cell line ADAR1 depletion RNA-Seq dataset was obtained from GEO (accession: GSE131658)(*18*). In brief, fastq files for scScramble, shADAR1#3, and shADAR1#9 were downloaded from SRA (accession: SRP199236) and aligned to the Ensembl hg38 reference genome using STAR (v2.7.11b)(*67*) with options for splice junction strand information (--outSAMstrandField intronMotif). Following alignment, intron retention events were identified using IRFinder (v2.0.0)(*20*) with the following stringent filtering criteria: no warnings; no known-exon overlap; coverage ≥ 0.9; intron depth ≥ 10; ΔIR ratio ≥ 0.05 or ≤ −0.05 in shADAR1 versus shScramble samples; and a consistent directional trend across both biological replicates.

### RNA sequencing and identification of A-to-I editing events

RNA-seq read alignment and transcript quantification were performed using RSEM (v1.3.3)(*68*) with STAR (v2.7.11b)(*67*) for alignment against the Ensembl GRCh38 human reference genome and gene annotation (Homo sapiens GRCh38, release 109) using default parameters. A-to-I RNA editing events were identified using REDItools3 and subsequently annotated for genomic features, common SNPs (avSNP150), and repetitive elements (RepeatMasker) using ANNOVAR(*69*). Known SNPs were excluded from further analysis, and only A-to-I editing sites residing within annotated Alu elements with a mean read coverage ≥20 and an editing frequency ≥1% were retained. Differential editing analysis between FLAG-ADAR1p110 versus empty vector (EV) and FLAG-ADAR1p90 versus EV conditions was conducted using DESeq2 (v1.46.0) (*70*) with the likelihood ratio test (LRT). Differentially edited sites were defined by a change in editing frequency (Δediting frequency) ≥5% and an adjusted *p*-value <0.1 (biological replicates, n = 3).

### WB analysis

Pelleted cells were lysed with RIPA buffer that is supplemented with 1× cOmplete EDTA-free protease inhibitor cocktail (Roche, COEDTAF-RO) and 1× PhosSTOP phosphatase inhibitor (Roche, PHOSS-RO). Protein lysates were clarified at 4°C for 10 minutes at 13,000 rpm on a table-top centrifuge. Protein concentrations were quantified using Bradford assay (Bio-Rad, 5000006). 20µg of protein lysates for each group were denatured in 1× Bolt LDS Sample Buffer (Invitrogen, B0007) for 70°C for 10 minutes prior to separation in a 4 – 8% Tris/Glycine SDS-PAGE gel for 1 hour 30 minutes at constant 100V prior to overnight wet transfer at 4°C using Tris/Glycine buffer with 20% methanol to a polyvinylidene difluoride (PVDF) membranes (Bio-Rad, 1620177). For SSO treatment samples, protein lysates were separated in a 4 – 6% SDS-PAGE gel for 1 hour 45 minutes. Transferred PVDF membranes were blocked with 5% (w/v) milk that is diluted in 1× TBS-T (0.1% (v/v) Tween- 20) for 1 hour at room temperature followed by 2 hours of primary antibody incubation or overnight at 4°C. Membranes were washed thrice with 1× TBS-T followed by incubation of secondary antibody for 1 hour at room temperature. Amersham enhanced chemiluminescence (ECL) (Cytiva, RPN2105) or SuperSignal™ West Femto (Thermo Fisher, 34096) was added to membranes and visualised using Chemidoc imaging system (Bio-Rad). If stripping of PVDF is required, especially the probing of total proteins after detection of phosphorylated proteins, membranes were stripped using WB stripping solution strong (Nacalai tesque, 05677-65) for 10 minutes at room temperature with agitation, followed by washing 3× with TBS-T.

Primary antibodies used in this study: anti-ADAR1 (Abcam, ab88574), anti-ADAR1 (Cell Signaling Technology, D7E2M), anti-β actin HRP (Santa Cruz Biotechnology, sc-47778 HRP), anti-FLAG (Proteintech, 20543-1-AP & 66008-4-Ig), anti-MYC-tag (Proteintech, 16286-1-AP & 60003-2-Ig), anti-GFP (Santa Cruz Biotechnology, sc-9996), anti-PKR (phosphor T446) (Abcam, ab32036), anti-PKR (Santa Cruz Biotechnology, sc-6282), anti-IRF3 (S386) (Abcam, ab76493), anti-IRF3 (Santa Cruz Biotechnology, sc-33641), anti-laminB1 (Proteintech, 12987-1-AP), anti-α tubulin (Santa Cruz Biotechnology, sc-32293), and anti-HNRNPA1 (Proteintech, 11176-1-AP). Secondary antibodies used in this study: HRP-linked anti-rabbit IgG (Cell Signaling Technology, 7074), HRP-linked anti-mouse IgG (Cell Signaling Technology, 7076).

### Splicing analysis by RT-qPCR or semiquantitative PCR

RNA was extracted using RNeasy mini kit (Qiagen) followed by DNaseI digestion on column. A total of 500 or 1 µg RNA was used for cDNA synthesis with the SensiFAST kit (Meridian Bioscience, BIO-65053). qPCR was carried out using GoTaq qPCR master mix (Promega, A600A) in a QuantStudio 5 system (Applied Biosystems, A34322) with a default comparative Ct protocol. Calculation of fold change (mRNA abundance) is represented as 2^−ΔΔCt^ where ΔCt = Ct_target_ – Ct_GAPDH_ and ΔΔCt = ΔCt_treatment_ – average ΔCt_control_. For the quantification of ADAR1 expression in CRC clinical samples (ΔΔCt), ΔCt_ADAR1_ = Ct_ADAR1_ – Ct_UBC_ and that ΔΔCt = ΔCt_sample_ – ΔCt_average normal_ whereby ΔCt_average normal_ is the average of all ΔCt_ADAR1_ of normal non-tumor samples. The calculation of splicing ratio by qPCR is in accordance with published protocols(*71*) in which ratio of splice isoforms is calculated as 2^−ΔCt^ whereby ΔCt = Ct_IR_ – Ct_non-IR_ and that relative fold change of splice ratios between experiment and control is represented as 2^−ΔCt(experiment)^/2^−ΔCt(control)^. Similarly, for the splicing ratio calculation in CRC samples, relative fold change of splice ratios between patient samples is represented as 2^−ΔCt(sample)^/2^−ΔCt(average^ ^normal)^. For semi-quantitative splicing analyses, RT- PCR was first performed using Platinum II Hot Start master mix (Invitrogen, 14001012) with cycling conditions in accordance with the manufacturer protocol. PCR products were separated using a 2% agarose gel (TBE) containing 1× GelRed Nucleic Acid Gel Stain (Merck, SCT123). Gel was imaged using Bio-rad GelDoc Go system and PCR bands semi-quantified using ImageJ. Percent-spliced-in (PSI) is defined as the band intensity of *ADAR1+I13* divided by the total abundance of *ADAR1+I13* and *ADAR1ΔI13*. Sequences of primers are listed in Supplementary Table 1.

### ADAR1 eCLIP-seq

The eCLIP was performed as per previously described(*28*). Briefly, anti-ADAR antibody (Sigma, HPA003890) or control IgG (Santa Cruz Biotechnology, sc-2025) was used to immunoprecipitate endogenous ADAR1 from parental EC109 cells. Next, the bound protein-RNA products were subjected to gel electrophoresis and membrane transfer. Bound RNAs on the membrane corresponding to the protein size of ADAR1 and 75 kDa above was extracted and further processed with adaptor ligation. The cDNA library was prepared by revere transcription and sequenced by paired-end 100 bp sequencing performed on the Illumina HiSeq 4000 platform. The sequencing data were processed as previously described and clusters identified in IP samples were compared against paired size-matched input to obtain significantly enriched peaks using a Fisher’s Exact test. Peaks with fold enrichment (4-fold) and significance (*p*-value <0.001) in immunoprecipitation versus paired size-matched input sample were defined as significant binding peaks.

### RNA immunoprecipitation (RIP)-quantitative PCR analysis

Harvested cell pellets were lysed in lysis buffer [50mM Tris/Cl pH7.5, 150mM NaCl, 0.5mM EDTA, 0.5% Igepal CA-630 (Sigma-Aldrich, 18896), supplemented with 1× cOmplete EDTA-free protease inhibitor cocktail (Roche, COEDTAF-RO) and SUPERase In RNase inhibitor (Invitrogen, AM2696)] for 30 minutes at 4°C. In assays that require treatment, cells were seeded in 10cm plates prior to transfection and subsequent harvesting with a cell scraper. Dynabeads protein G (Invitrogen, 10004D) were equilibrated in dilution buffer [50mM Tris/Cl pH7.5, 150mM NaCl, 0.5mM EDTA, 0.05% Igepal CA-630 (Sigma-Aldrich, 18896)] prior to aliquoting (20uL bead slurry per reaction) for antibody conjugation: Rabbit IgG isotype control control (Novus Biologicals, NB810-56910), FLAG (Proteintech, 20543-1-AP), or hnRNPA1 (Proteintech, 11176-1-AP) for 1 hour at 4°C with rotation. Conjugated beads were incubated overnight with cell lysates at 4°C with rotation. Immunoprecipitates were washed thrice using the dilution buffer prior to separation of immunoprecipitates - 10% for protein enrichment validation via western blot, 90% for RNA extraction using RNeasy mini kit (Qiagen, 74104). Extracted RNA subsequent underwent first-strand cDNA synthesis using SensiFAST kit (Meridian Bioscience, BIO-65053), followed by qPCR analysis. %Input = 2 ^−ΔCt^ × 100%; ΔCt = Ct_RIP_ – [Ct_input_ – dilution factor]. Sequences of primers are listed in Supplementary Table 1.

### Rapid Amplification of cDNA Ends (RACE)

5’- and 3’-RACE ready cDNA was generated from HEK293T total RNA using the SMARTer® RACE 5’/3’ Kit (Takara, 634858). 5’- and 3’-gene specific primers (GSPs) were designed to target intron 13 of the *ADAR1* transcript for 5’RACE and 3’RACE, respectively. 5’ and 3’RACE PCR was performed using Platinum SuperFi II Green PCR Master Mix (Invitrogen, 12369010). 5’ and 3’RACE PCR products were subsequently separated using a 1% agarose gel (Tris/Borate/EDTA buffer) containing 1× GelRed® Nucleic Acid Gel Stain (Merck, SCT123). PCR bands between ∼3 to 4kbp were gel-purified using QIAquick Gel Extraction kit (Qiagen, 28704) according to the instructions of the manufacturer. In-Fusion cloning of purified PCR products into linearised pRACE vector was performed and subsequently transformed into Stellar competent cells (Takara, 636763) according to the manufacturer protocol. Ten colonies were identified and processed for Sanger sequencing using multiple sequencing primers targeting different regions of the *ADAR1* transcript and subsequently visualized using SnapGene Viewer. Sequences of primers are listed in Supplementary Table 1.

### Peptide-enrichment analysis from immunoprecipitation followed by mass spectrometry (IP-MS) data

Peptide and protein identifications from our previously published ADAR1 IP-MS dataset (27) were validated using Scaffold software (Proteome Software, Inc.). Raw MS/MS spectra were searched against the appropriate protein database and subsequently imported into Scaffold, where identifications were filtered using the following criteria: a minimum protein probability of 95%, a minimum peptide probability of 90%, and a minimum of two unique peptides per protein. Quantitative analysis was performed by spectral counting, and fold-change enrichment between immunoprecipitated samples and IgG control samples was calculated to identify high-confidence interacting proteins. Enriched proteins were further filtered to retain only those showing greater than 5% enrichment over the IgG control, and non-RNA-binding proteins (non-RBPs) were subsequently excluded to focus the analysis on biologically relevant interaction candidates.

### Co-immunoprecipitation (Co-IP)

HEK293T cells were seeded in 6-well plates for 24 hours prior to transfection for 24- or 48- hours. Cell pellets were harvested and lysed in co-IP lysis buffer [10mM Tris/Cl pH7.5, 150mM NaCl, 0.5mM EDTA, 0.5% Igepal CA-630 (Sigma-Aldrich, 18896), supplemented with 1× cOmplete™ EDTA-free protease inhibitor cocktail (Roche, COEDTAF-RO)] for 30 minutes at 4°C. Protein lysates were clarified at max speed at 4°C for 10 minutes and subsequently quantified using Bradford assay (Bio-Rad, 5000006). 400µg of cell lysates were diluted in 1mL of dilution buffer (10mM Tris/Cl pH7.5, 150mM NaCl, 0.5mM EDTA). 5% volume from 1mL was used as input. For FLAG-tag IP, 15µL of Fab-Trap™ Agarose (Chromotek®, ffa) beads were utilised for each reaction. IP was performed for 1 hour prior to washing 3× using wash buffer [(10mM Tris/Cl pH7.5, 150mM NaCl, 0.5mM EDTA, 0.05% Igepal CA-630 (Sigma-Aldrich, 18896)]. Subsequent protein denaturation and SDS-PAGE are described as in 2.10, with a final input of 1.25% relative to IP.

### Actinomycin D Transcriptional Inhibition Assay

HEK293T cells were seeded in 6-well plates 24 hours prior to treatment. Transcription was globally inhibited by treatment with actinomycin D (ActD; Sigma, A1410) at a final concentration of 1 µg/mL, or an equivalent volume of DMSO as a vehicle control. Total RNA was harvested at 0, 2, and 4 hours post-treatment using standard RNA extraction methods. The abundance of intron 13-retained (+I13) and intron 13-skipped (ΔI13) ADAR1 transcripts was quantified by RT-qPCR using isoform-specific primers. mRNA abundance at each timepoint was normalized to the 0-hour timepoint and expressed as fold change, and transcript decay rates were compared between +I13 and ΔI13 isoforms.

### Nascent RNA Capture Assay (Click-iT™ RNA)

To directly measure the stability of nascent ADAR1 transcripts, the Click-iT™ RNA Imaging Kit (Invitrogen, C10365) was performed according to the manufacturer’s instructions. Briefly, HEK293T cells were seeded in 6-well plates and pulse-labeled with 5-ethynyl uridine (EU) for 24 hours to allow EU incorporation into newly synthesized RNA. Following pulse labeling, cells were washed and incubated in EU-free medium for a chase period of 0 or 6 hours to track RNA decay over time. EU-labeled nascent RNA was subsequently isolated by click chemistry-based biotinylation and captured using streptavidin-coated magnetic beads. The abundance of intron 13-retained (+I13) and intron 13-skipped (ΔI13) ADAR1 transcripts within the captured nascent RNA fraction was quantified by RT-qPCR using isoform-specific primers. Fold loss of RNA between the 0- and 6-hour chase timepoints was calculated and compared between +I13 and ΔI13 isoforms to assess their relative stability.

### Cell fractionation assays

HEK293T cells were transfected with ADAR1 isoform overexpression constructs for 48 hours prior to acquiring nuclear and cytoplasmic fractions using the Nuclear Extract kit (Active Motif, 40010) according to the manufacturer protocol. For assays without treatment, cells were seeded for at least 48 hours prior to fractionation. Alterations of the extraction kit include 5minute incubation with 1× hypotonic buffer instead of 15 minutes for the acquisition of cytoplasmic fraction. Prior to nuclear fraction lysis, nuclear pellet was washed 3 times using 1×PBS. Nuclear pellet and whole cell lysate fractions were lysed using 1× RIPA (Radioimmunoprecipitation assay) (Sigma, R0278) buffer supplemented with 1× cOmplete EDTA-free protease inhibitor cocktail (Roche, COEDTAF-RO). HRP-conjugated α-tubulin (Proteintech, HRP-66031) and lamin-B1 (Proteintech, 12987-1-AP) were used as cytoplasmic and nuclear markers, respectively.

### Immunofluorescence (IF) Staining

Cells were cultured on coverslips for 24 hours. Cells were washed with 1 × phosphate-buffered saline (PBS; 10 mM phosphate, 137 mM NaCl and 2.7 mM potassium chloride (KCl)) before fixation with 4% paraformaldehyde solution for 10 min at RT. Fixed cells were washed with 1 × PBS thrice, at every 5 min interval. Subsequently, cells were permeabilized with 0.5% Triton-X100 in 1 × PBS for 10 min at RT, followed by three washes with 1 × PBS at every 5 min interval. Serum-free protein block (Dako) was used and cells were blocked for 1 h at RT. Cells were blotted for 1 h at RT, with the following primary antibodies: rabbit anti-ADAR1 (1:200; CST #14175), mouse anti-α-tubulin (1:200; Santa Cruz, sc-32293). Cells were washed thrice with 0.1% Tween-20 in 1 × PBS (PBST). Cells were co-stained in the dark with 4′,6-diamidino-2-phenylindole (DAPI; 0.2 µg/ml) and the following secondary antibodies for 30 min at RT: Coralite594-conjugated Goat Anti-Rabbit IgG (1:1000; Proteintech, #SA00013-4) and Coralite488-conjugated Goat Anti-Mouse IgG (1:1000; Proteintech, SA00013-1); followed by three washes with PBST. The immunostained cells on the coverslips were mounted onto slides using SlowFade Gold antifade mountant (ThermoFisher Scientific). Immunostained cells were viewed using Olympus FV1000 confocal microscope.

### Biotin-labelled RNA pulldown assay

DNA templates for *in vitro* transcription were generated by PCR amplification or primer extension using primers containing a T7 promoter sequence (5′-CGAAATTAATACGACTCACTATAGGG-3′) at the forward primer. Biotin-labelled RNA probes were synthesised using the T7 RiboMAX™ Large Scale RNA Production System (Promega) in the presence of biotin-UTP, according to the manufacturer’s instructions. Newly synthesised RNA was purified using the RNeasy Mini Kit (Qiagen, 74104) or the mirVana miRNA Isolation Kit (Ambion) and assessed for integrity prior to use.

For pulldown assays, HEK293T cells overexpressing MYC-tagged ADAR1p110, ADAR1p90, or EV were lysed in lysis buffer [50 mM Tris-HCl pH 7.5, 150 mM NaCl, 0.5 mM EDTA, 0.5% Igepal CA-630 (Sigma-Aldrich, 18896), supplemented with 1× cOmplete EDTA-free protease inhibitor cocktail (Roche, COEDTAF-RO) and SUPERase In RNase inhibitor (Invitrogen, AM2696)] for 30 minutes at 4°C. Clarified lysates were pre-cleared with Dynabeads MyOne Streptavidin C1 beads (Invitrogen) for 30 minutes at 4°C with rotation to remove non-specific binding. Biotin-labelled RNA probes were denatured at 95°C for 2 minutes and allowed to re-fold at room temperature for 20 minutes prior to incubation with pre-cleared lysates overnight at 4°C with rotation. Streptavidin beads equilibrated in dilution buffer [50 mM Tris-HCl pH 7.5, 150 mM NaCl, 0.5 mM EDTA, 0.05% Igepal CA-630 (Sigma-Aldrich, 18896)] were added and incubated for 1 hour at 4°C with rotation to capture biotin-RNA-protein complexes. Beads were washed three times with dilution buffer, and bound proteins were eluted by boiling in SDS sample buffer and resolved by SDS-PAGE for western blot analysis with antibodies against PKR and ADAR1.

### Cytoplasmic J2 (anti-dsRNA) eCLIP-seq

Cytoplasmic dsRNA-eCLIP sequencing was modified from standard eCLIP sequencing(*72*). Cytoplasm of the same number of cells from a 15 cm dish was fractionated using 500 μL PARIS cell fraction buffer, topped up to 1.5mL with lysis buffer (50 mM Tris pH 7.4, 100 mM NaCl, 1% NP40, 0.1% SDS, 0.5% sodium deoxycholate, protease inhibitor) and precleared with Dynabeads™ Protein G (Invitrogen, 10004D) coated with mouse IgG (Santa Cruz Biotechnology, sc-2025). Samples were incubated with 5 μg J2 dsRNA antibody or mouse IgG, UV crosslinked (400 mJ/cm², 254 nm), sonicated (Bioruptor on the low setting in 4°C for 5 min, cycling 30 seconds on and 30 seconds off), and treated with TURBO™ DNase (Invitrogen, AM2238) and Ambion™ RNase I (Invitrogen, AM2295). RNA-protein complexes were immunoprecipitated using Dynabeads™ Protein G, washed with high salt buffer (50 mM Tris pH 7.4, 1 M NaCl, 1 mM EDTA, 1% NP40, 0.1% SDS, 0.5% sodium deoxycholate) and wash buffer (20 mM Tris pH 7.4, 10 mM MgCl2, 0.2% Tween-20), and digested with Proteinase K. RNA was extracted using TRIzol™ (Invitrogen, 15596018) and Zymo RNA Clean & Concentrator Kit (Zymo Research, R1019), dephosphorylated, adapter-ligated, reverse-transcribed, and sequenced on BGI DNBseq-G400. Two independent eCLIP were performed for each sample.

eCLIP-Seq read files were pre-processed and analysed using the eCLIP-seq CLIPper peak enrichment pipeline(*28*) with hg19 (Gencode GRCh37). Peak enrichment scores for each CLIP samples were normalised using the corresponding input samples. Enriched peaks were defined as peaks with log_2_(Fold change) > 1 and log_10_(*P*-value) < 2. Functional and repeat element annotation of enriched peaks in the HEK293T was performed using HOMER annotatePeaks using the hg19 genome annotation(*73*). For differential peak analysis, enriched peaks from all samples were concatenated and merged for read depth analysis using samtools bedcov (v1.5) and imported into R for subsequent analyses. To account for potential systematic enrichment in high complexity genomic regions and library size effects, read counts of CLIP samples were subtracted by counts in the corresponding input samples at each peak and normalised using library size factors calculated from the input samples(*74*). Differential binding analysis was performed using the DESeq2 R package (v1.42.1)(*70*). The statistical significance threshold for differential binding was set at absolute log_2_Fold change > 1 and BH-adjusted *P*-value < 0.01.

### Clinical CRC sample cohort

All human tissue samples used in this study were obtained with informed consent and approved by the Domain-Specific Review Board (DSRB) under the National Healthcare Group (NHG), Singapore. For RT-qPCR analysis of *ADAR1* transcript expression and intron 13 retention in clinical specimens, 32 matched CRC tumour and adjacent non-tumour (NT) pairs were selected from the cohort based on comparable expression of the housekeeping gene *UBC* between matched samples (cycle threshold [C_t_] difference ≤1 cycle), ensuring reliable normalisation across paired samples.

### *In vivo* tumorigenicity assay

To test the effect of ADAR1p110 or ADAR1p90 overexpression in tumorigenesis, 1×10^6^ EV or p110-or p90-overexpressing HCT116 cells were subcutaneously injected into the left and right flanks of 6 to 8 weeks old NSG mice. At end point, tumors were harvested for downstream analyses. All animal experiments were approved by the Institutional Animal Care and Use Committees (IACUC) of National University of Singapore (NUS; Singapore) with the protocol number R21-1350. NSG mice were 6 to 8 weeks old and were acquired through breeding protocol BR22-0577 and housed under IACUC mice housing policies.

## SUPPLEMENTARY FIGURE LEGENDS

**Fig. S1.**
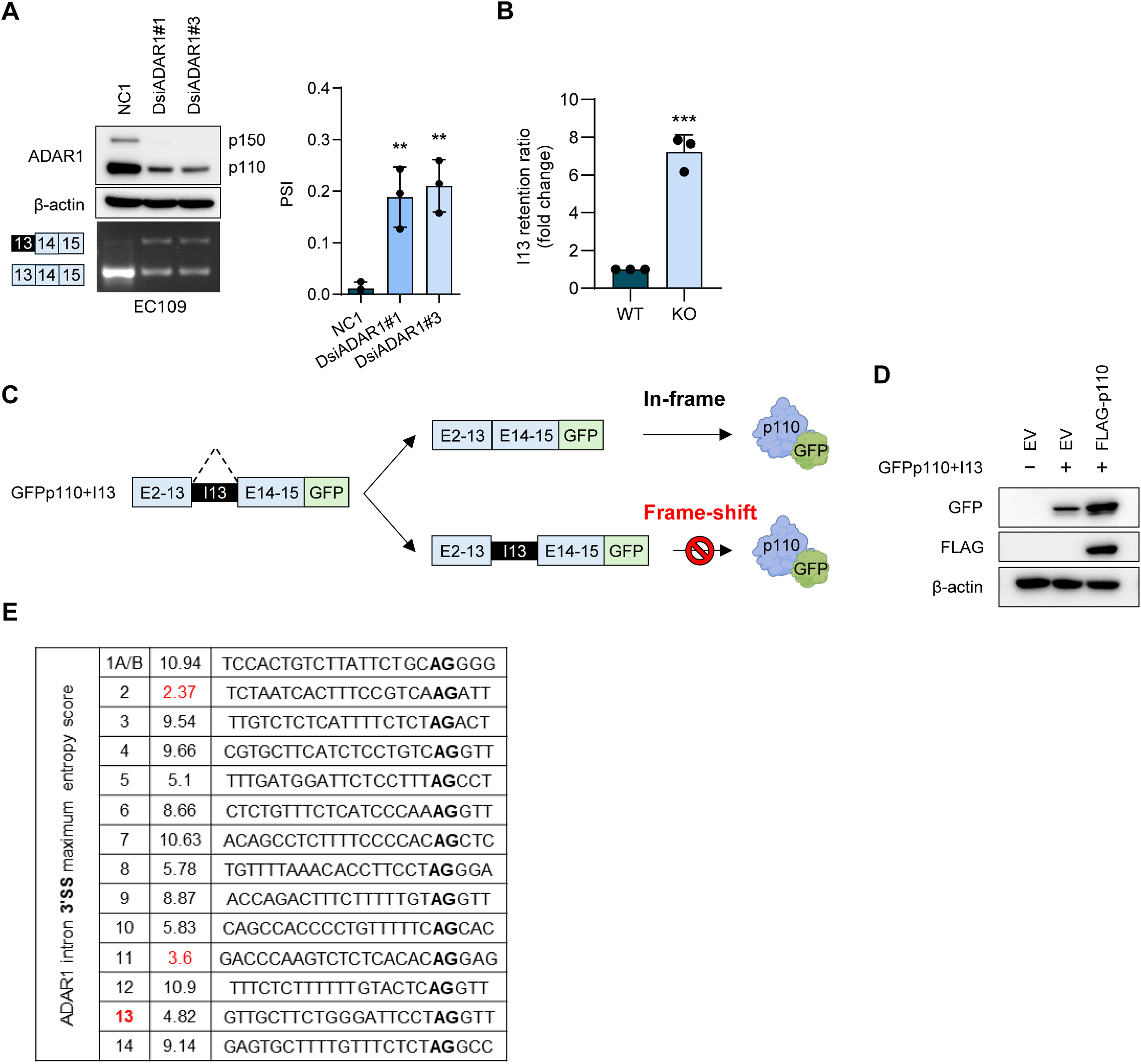
ADAR1 negatively regulates retention of intron 13 in its own transcripts. **(A)** Left, top: WB analysis of ADAR1 expression in ADAR1 KD and NC1 EC109 cells, with β-actin as the loading control. Left, bottom: RT-PCR detection of intron 13-retained and intron 13-excised transcripts in the same groups. Right: Bar charts showing PSI values for the same groups. **(B)** qPCR quantification of I13R ratios (fold change) in ADAR1-knockout (KO) HEK293T cells relative to wild-type (WT) HEK293T cells. Fold change is calculated by normalizing the I13R ratio in KO cells to that in WT cells. **(C)** Schematic of the GFP fusion reporter (GFPp110+I13), in which the full ADAR1 p110 coding sequence with intron 13 inserted between exons 13 and 14 is placed upstream of the GFP open reading frame (ORF). **(D)** WB analysis of the indicated proteins in HEK293T cells co-transfected with FLAG-p110 or empty vector (EV) and the GFPp110+I13 reporter plasmid. **(E)** *In silico* maximum entropy scores for the indicated ADAR1 intronic 3′SS. Scores lower than that of the intron 13 3′SS are highlighted in red. The 3′SS dinucleotide AG is shown in bold. **(A, B)** Data are presented as mean ± SD (n = 3 biological replicates). Each data point represents a biological replicate. Statistical significance is determined by two-tailed Student’s *t*-test (paired for (**A**); unpaired for (**B**)) (** *p* < 0.01; *** *p* < 0.001).

**Fig. S2.**
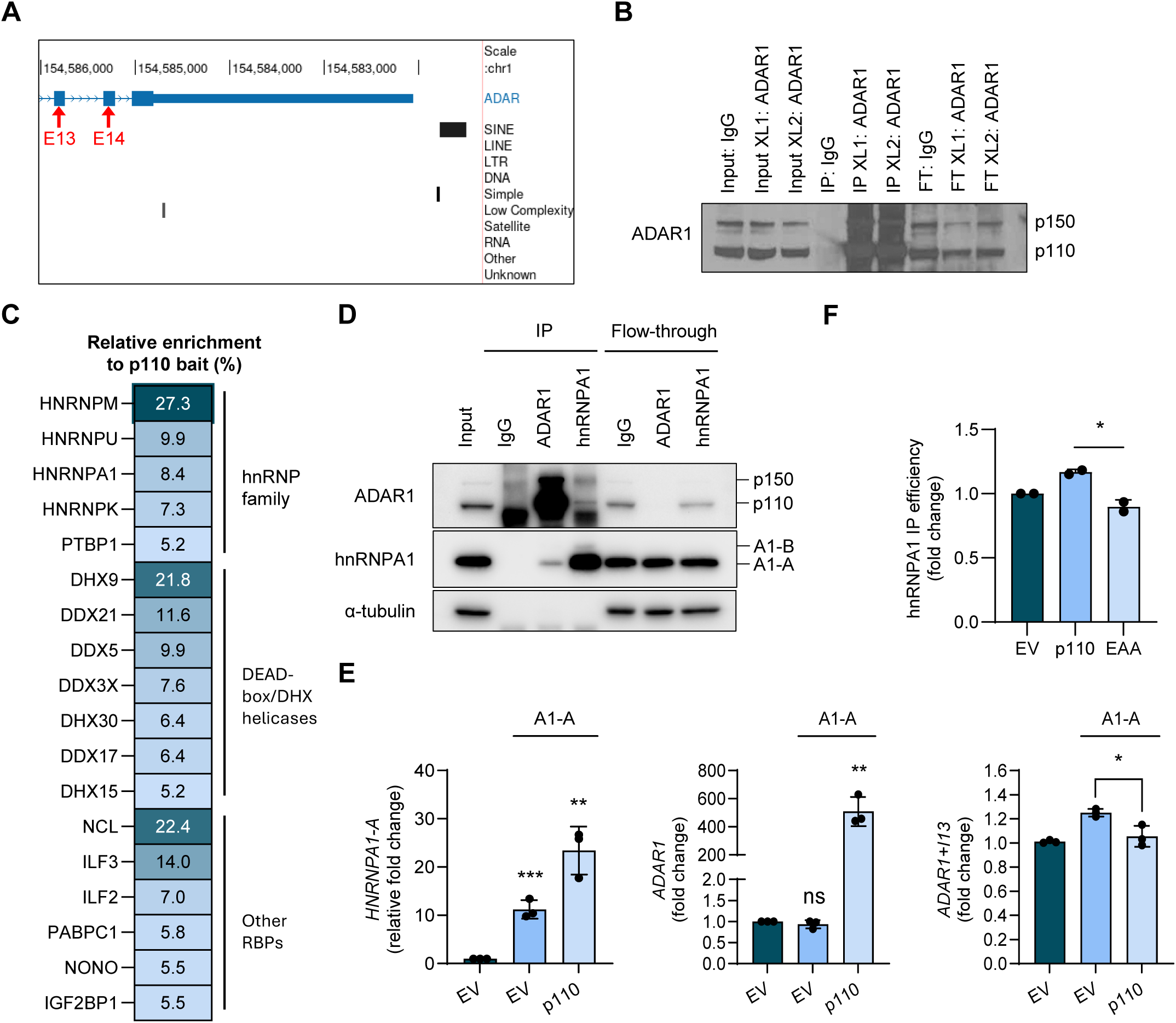
ADAR1 binds its own exons 13 and 14 and counteracts hnRNPA1 association with intron 13 to prevent retention. **(A)** UCSC Genome Browser snapshot of the *ADAR1* region spanning exon 13 to exon 15, with RepeatMasker-annotated elements indicated (black bars). Exons 13 and 14 are indicated with red arrows. **(B)** WB analysis of ADAR1 expression levels in the indicated ADAR1 eCLIP samples. Input corresponds to 2% of total cell lysate. FT, flow-through. **(C)** Relative peptide enrichment with the ADAR1p110 bait from our previously published ADAR1p110 IP-MS dataset, filtered for >5% enrichment and excluding non-RNA-binding proteins (non-RBPs). **(D)** WB analysis of ADAR1, hnRNPA1 (including the alternatively spliced isoforms A1-A and A1-B) and α-tubulin in ADAR1, hnRNPA1 or IgG immunoprecipitates in HEK293T cells. Input corresponds to 5% of total cell lysate. **(E)** RT-qPCR analysis of *hnRNPA1-A*, *ADAR1* and endogenous *ADAR1+I13* transcript levels in HEK293T cells transfected with EV alone or co-transfected with hnRNPA1-A and EV or ADAR1p110. **(F)** Quantification of hnRNPA1 band intensity in the samples shown in **Fig. 3D**. Band intensities were measured using ImageJ and normalized to the EV control. **(E, F)** Data are presented as mean ± SD (n = 3 biological replicates for (**E)**; n = 2 biological replicates for (**F**)). Each data point represents a biological replicate. Statistical significance is determined by two-tailed, unpaired Student’s *t*-test (* *p* < 0.05; ** *p* < 0.01; *** *p* < 0.001; *ns*, not significant).

**Fig. S3.**
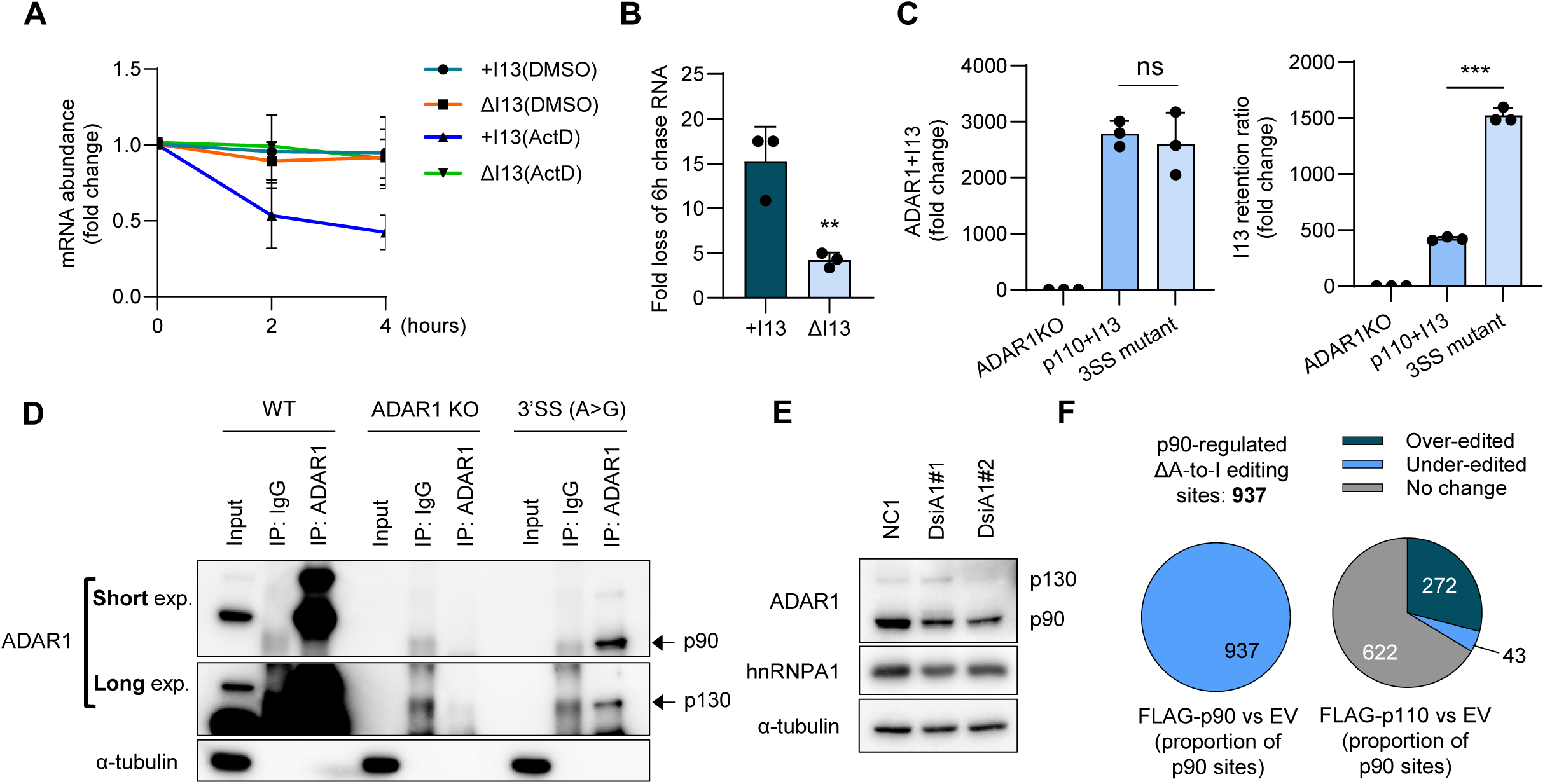
ADAR1p90 is encoded by a less stable intron 13-retained transcript, the levels of which are reduced by hnRNPA1 knockdown. **(A)** RT-qPCR analysis of intron 13-retained (+I13) and intron 13-excised (ΔI13) ADAR1 transcript abundance in HEK293T cells treated with actinomycin D (ActD) or DMSO vehicle control for the indicated durations (0, 2, and 4 hours). mRNA abundance is expressed as fold change relative to the 0-hour timepoint. Data are presented as mean ± SD from three biological replicates. **(B)** RT-qPCR analysis of nascent RNA stability using the Click-iT EU assay. HEK293T cells were pulsed with 5-ethynyl uridine (EU) for 24 hours, followed by a 6-hour EU-free chase. Data are shown as fold loss of EU-labelled +I13 and ΔI13 transcripts after the chase period. **(C)** RT-qPCR analysis of intron 13-retained *ADAR1* transcript levels (left) and the I13R ratio (right) in ADAR1-KO HEK293T cells or ADAR1-KO cells transduced with the ADAR1p110+I13 or 3′SS mutant constructs described in **Fig. 4C**. **(D)** WB analysis of ADAR1 isoforms and α-tubulin in ADAR1 immunoprecipitates (IP: ADAR1) from WT, ADAR1-KO, and 3′SS mutant HEK293T cells. IgG antibody serves as a negative control, and input corresponds to 5% of total cell lysate. **(E)** WB analysis of ADAR1 and hnRNPA1 protein levels in 3′SS mutant cells treated with a negative control siRNA (NC1) or two independent DsiRNA targeting *HNRNPA1* (DsiA#1 and DsiA#2). α-tubulin serves as a loading control. Blot shown is representative of 2 independent biological replicates. **(F)** Pie charts showing the classification of A-to-I editing changes at the 937 high-confidence ADAR1p90-regulated sites identified by RNA-seq. Sites are categorised as over-edited, under-edited, or no change relative to EV control. Left: editing status of the 937 p90-regulated sites in FLAG-ADAR1p90-overexpressing cells. Right: editing behaviour of the same 937 sites upon FLAG-ADAR1p110 overexpression, using identical filter criteria. (**B, C**) Data are presented as mean ± SD (n = 3 biological replicates); statistical significance is determined by two-tailed, unpaired Student’s *t*-test (** *p*< 0.01; *** *p*< 0.001; *ns*, not significant).

**Fig. S4.**
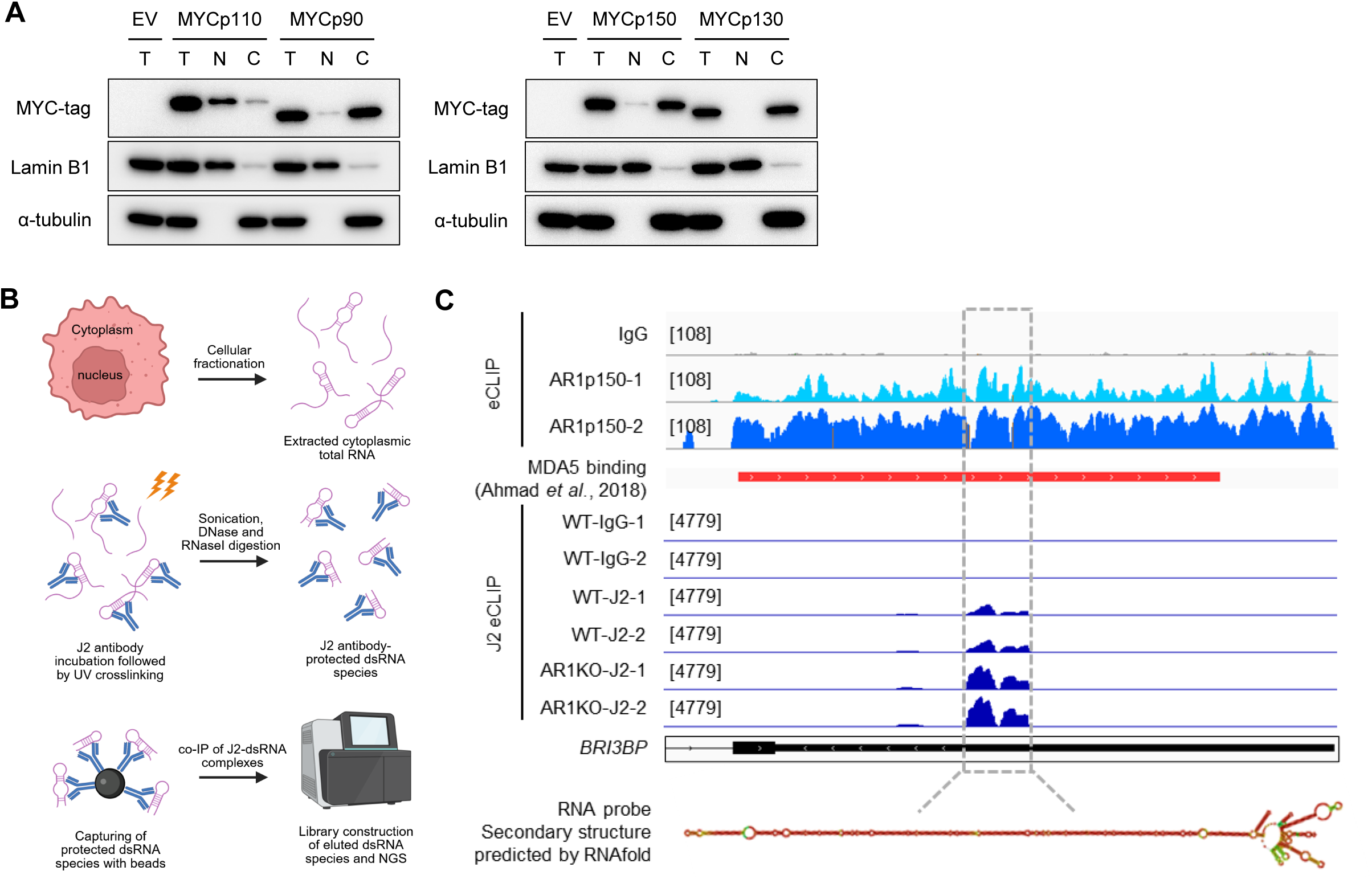
Cytoplasmic localisation of ADAR1p90 and identification of the *BRI3BP* immunogenic dsRNA substrate. **(A)** WB analysis shows fractionation into Total (T), Nuclear (N), and Cytoplasmic (C) for EV, MYCp110, and MYCp90 (left blot), and EV, MYCp150, and MYCp130 (right blot). Lamin B1 is enriched in N fractions and α-tubulin in C fractions, confirming clean fractionation. **(B)** Schematic of in-house developed cytoplasmic J2-eCLIP-seq pipeline. **(C)** IGV tracks across the *BRI3BP* 3’UTR showing eCLIP-seq binding peaks for ADAR1p150 and J2 (dsRNA) from our in-house ADAR1p150 eCLIP-seq and cytoplasmic J2 eCLIP-seq datasets. The red bar marks peaks identified by a published MDA5 protection assay that maps transcriptome-wide MDA5 binding sites. Bottom: in silico prediction of RNA secondary structure within the boxed region using RNAfold. Minimum free energy (MFE) structures with base-pairing probabilities are shown.

## REFERENCES

1. C. X. Chen et al., A third member of the RNA-specific adenosine deaminase gene family, ADAR3, contains both single- and double-stranded RNA binding domains. RNA 6, 755–767 (2000).

2. U. Kim, Y. Wang, T. Sanford, Y. Zeng, K. Nishikura, Molecular cloning of cDNA for double-stranded RNA adenosine deaminase, a candidate enzyme for nuclear RNA editing. Proc Natl Acad Sci U S A 91, 11457–11461 (1994).

3. T. Melcher et al., RED2, a brain-specific member of the RNA-specific adenosine deaminase family. J Biol Chem 271, 31795–31798 (1996).

4. T. Melcher et al., A mammalian RNA editing enzyme. Nature 379, 460–464 (1996).

5. M. F. Schneider, J. Wettengel, P. C. Hoffmann, T. Stafforst, Optimal guideRNAs for re-directing deaminase activity of hADAR1 and hADAR2 in trans. Nucleic Acids Res 42, e87 (2014).

6. H. Chung et al., Human ADAR1 Prevents Endogenous RNA from Triggering Translational Shutdown. Cell 172, 811–824 e814 (2018).

7. H. S. Gannon et al., Identification of ADAR1 adenosine deaminase dependency in a subset of cancer cells. Nat Commun 9, 5450 (2018).

8. S. B. Hu et al., ADAR1p150 prevents MDA5 and PKR activation via distinct mechanisms to avert fatal autoinflammation. Mol Cell 83, 3869–3884 e3867 (2023).

9. K. Sinigaglia et al., An ADAR1 dsRBD3-PKR kinase domain interaction on dsRNA inhibits PKR activation. Cell Rep 43, 114618 (2024).

10. C. X. George, C. E. Samuel, Human RNA-specific adenosine deaminase ADAR1 transcripts possess alternative exon 1 structures that initiate from different promoters, one constitutively active and the other interferon inducible. Proc Natl Acad Sci U S A 96, 4621–4626 (1999).

11. J. B. Patterson, C. E. Samuel, Expression and regulation by interferon of a double-stranded-RNA-specific adenosine deaminase from human cells: evidence for two forms of the deaminase. Mol Cell Biol 15, 5376–5388 (1995).

12. J. M. Desterro et al., Dynamic association of RNA-editing enzymes with the nucleolus. J Cell Sci 116, 1805–1818 (2003).

13. J. Fritz et al., RNA-regulated interaction of transportin-1 and exportin-5 with the double-stranded RNA-binding domain regulates nucleocytoplasmic shuttling of ADAR1. Mol Cell Biol 29, 1487–1497 (2009).

14. K. Nishikura, A-to-I editing of coding and non-coding RNAs by ADARs. Nat Rev Mol Cell Biol 17, 83–96 (2016).

15. A. Strehblow, M. Hallegger, M. F. Jantsch, Nucleocytoplasmic distribution of human RNA-editing enzyme ADAR1 is modulated by double-stranded RNA-binding domains, a leucine-rich export signal, and a putative dimerization domain. Mol Biol Cell 13, 3822–3835 (2002).

16. K. Pakos-Zebrucka et al., The integrated stress response. EMBO Rep 17, 1374–1395 (2016).

17. J. J. Ishizuka et al., Loss of ADAR1 in tumours overcomes resistance to immune checkpoint blockade. Nature 565, 43–48 (2019).

18. S. J. Tang et al., Cis- and trans-regulations of pre-mRNA splicing by RNA editing enzymes influence cancer development. Nat Commun 11, 799 (2020).

19. C. Lorenzi et al., IRFinder-S: a comprehensive suite to discover and explore intron retention. Genome Biol 22, 307 (2021).

20. R. Middleton et al., IRFinder: assessing the impact of intron retention on mammalian gene expression. Genome Biol 18, 51 (2017).

21. J. J. Wong et al., Orchestrated intron retention regulates normal granulocyte differentiation. Cell 154, 583–595 (2013).

22. G. Yeo, C. B. Burge, Maximum entropy modeling of short sequence motifs with applications to RNA splicing signals. J Comput Biol 11, 377–394 (2004).

23. K. S. Pollard, M. J. Hubisz, K. R. Rosenbloom, A. Siepel, Detection of nonneutral substitution rates on mammalian phylogenies. Genome Res 20, 110–121 (2010).

24. A. Siepel et al., Evolutionarily conserved elements in vertebrate, insect, worm, and yeast genomes. Genome Res 15, 1034–1050 (2005).

25. A. Siepel, D. Haussler, Combining phylogenetic and hidden Markov models in biosequence analysis. J Comput Biol 11, 413–428 (2004).

26. R. Lorenz et al., ViennaRNA Package 2.0. Algorithms Mol Biol 6, 26 (2011).

27. H. Hong et al., Bidirectional regulation of adenosine-to-inosine (A-to-I) RNA editing by DEAH box helicase 9 (DHX9) in cancer. Nucleic Acids Res 46, 7953–7969 (2018).

28. E. L. Van Nostrand et al., Robust transcriptome-wide discovery of RNA-binding protein binding sites with enhanced CLIP (eCLIP). Nat Methods 13, 508–514 (2016).

29. B. J. Liddicoat et al., RNA editing by ADAR1 prevents MDA5 sensing of endogenous dsRNA as nonself. Science 349, 1115–1120 (2015).

30. T. Sun et al., ADAR1 editing is necessary for only a small subset of cytosolic dsRNAs to evade MDA5-mediated autoimmunity. Nat Genet 57, 3101–3111 (2025).

31. P. Barraud, S. Banerjee, W. I. Mohamed, M. F. Jantsch, F. H. Allain, A bimodular nuclear localization signal assembled via an extended double-stranded RNA-binding domain acts as an RNA-sensing signal for transportin 1. Proc Natl Acad Sci U S A 111, E1852–1861 (2014).

32. S. Ahmad et al., Breaching Self-Tolerance to Alu Duplex RNA Underlies MDA5-Mediated Inflammation. Cell 172, 797–810 e713 (2018).

33. S. M. Rueter, T. R. Dawson, R. B. Emeson, Regulation of alternative splicing by RNA editing. Nature 399, 75–80 (1999).

34. X. Roca et al., Widespread recognition of 5’ splice sites by noncanonical base-pairing to U1 snRNA involving bulged nucleotides. Genes Dev 26, 1098–1109 (2012).

35. X. Roca, A. R. Krainer, Recognition of atypical 5’ splice sites by shifted base-pairing to U1 snRNA. Nat Struct Mol Biol 16, 176–182 (2009).

36. X. Roca, A. R. Krainer, I. C. Eperon, Pick one, but be quick: 5’ splice sites and the problems of too many choices. Genes Dev 27, 129–144 (2013).

37. L. Song et al., microRNA-451-modulated hnRNP A1 takes a part in granulocytic differentiation regulation and acute myeloid leukemia. Oncotarget 8, 55453–55466 (2017).

38. Y. Chen et al., Cooperative regulation of Zhx1 and hnRNPA1 drives the cardiac progenitor-specific transcriptional activation during cardiomyocyte differentiation. Cell Death Discov 9, 244 (2023).

39. M. Sukma, M. Tohda, H. Watanabe, K. Matsumoto, The mRNA expression differences of RNA editing enzymes in differentiated and undifferentiated NG108-15 cells. J Pharmacol Sci 98, 467–470 (2005).

40. C. Anadon et al., Gene amplification-associated overexpression of the RNA editing enzyme ADAR1 enhances human lung tumorigenesis. Oncogene 35, 4422 (2016).

41. A. R. Baker, F. J. Slack, ADAR1 and its implications in cancer development and treatment. Trends Genet 38, 821–830 (2022).

42. T. H. Chan et al., A disrupted RNA editing balance mediated by ADARs (Adenosine DeAminases that act on RNA) in human hepatocellular carcinoma. Gut 63, 832–843 (2014).

43. T. H. Chan et al., ADAR-Mediated RNA Editing Predicts Progression and Prognosis of Gastric Cancer. Gastroenterology 151, 637–650 e610 (2016).

44. L. Chen et al., Recoding RNA editing of AZIN1 predisposes to hepatocellular carcinoma. Nat Med 19, 209–216 (2013).

45. Y. Chen, H. Wang, W. Lin, P. Shuai, ADAR1 overexpression is associated with cervical cancer progression and angiogenesis. Diagn Pathol 12, 12 (2017).

46. Q. Jiang et al., ADAR1 promotes malignant progenitor reprogramming in chronic myeloid leukemia. Proc Natl Acad Sci U S A 110, 1041–1046 (2013).

47. E. Lazzari et al., Alu-dependent RNA editing of GLI1 promotes malignant regeneration in multiple myeloma. Nat Commun 8, 1922 (2017).

48. X. Liu et al., ADAR1 promotes the epithelial-to-mesenchymal transition and stem-like cell phenotype of oral cancer by facilitating oncogenic microRNA maturation. J Exp Clin Cancer Res 38, 315 (2019).

49. N. Paz-Yaacov et al., Elevated RNA Editing Activity Is a Major Contributor to Transcriptomic Diversity in Tumors. Cell Rep 13, 267–276 (2015).

50. Y. R. Qin et al., Adenosine-to-inosine RNA editing mediated by ADARs in esophageal squamous cell carcinoma. Cancer Res 74, 840–851 (2014).

51. J. Ramirez-Moya, A. R. Baker, F. J. Slack, P. Santisteban, ADAR1-mediated RNA editing is a novel oncogenic process in thyroid cancer and regulates miR-200 activity. Oncogene 39, 3738–3753 (2020).

52. K. Shigeyasu, et al., AZIN1 RNA editing confers cancer stemness and enhances oncogenic potential in colorectal cancer. JCI Insight 3, (2018).

53. Y. Sun et al., The aberrant expression of ADAR1 promotes resistance to BET inhibitors in pancreatic cancer by stabilizing c-Myc. Am J Cancer Res 10, 148–163 (2020).

54. H. Sato, R. H. Singer, Cellular variability of nonsense-mediated mRNA decay. Nat Commun 12, 7203 (2021).

55. M. C. Dyle, D. Kolakada, M. A. Cortazar, S. Jagannathan, How to get away with nonsense: Mechanisms and consequences of escape from nonsense-mediated RNA decay. Wiley Interdiscip Rev RNA 11, e1560 (2020).

56. R. Karam et al., The unfolded protein response is shaped by the NMD pathway. EMBO Rep 16, 599–609 (2015).

57. Z. Li, J. K. Vuong, M. Zhang, C. Stork, S. Zheng, Inhibition of nonsense-mediated RNA decay by ER stress. RNA 23, 378–394 (2017).

58. Y. S. Oren et al., The unfolded protein response affects readthrough of premature termination codons. EMBO Mol Med 6, 685–701 (2014).

59. K. Degenhardt et al., Autophagy promotes tumor cell survival and restricts necrosis, inflammation, and tumorigenesis. Cancer Cell 10, 51–64 (2006).

60. L. B. Gardner, Hypoxic inhibition of nonsense-mediated RNA decay regulates gene expression and the integrated stress response. Mol Cell Biol 28, 3729–3741 (2008).

61. C. Lior et al., Mapping the tumor stress network reveals dynamic shifts in the stromal oxidative stress response. Cell Rep 43, 114236 (2024).

62. M. W. Popp, L. E. Maquat, Nonsense-mediated mRNA Decay and Cancer. Curr Opin Genet Dev 48, 44–50 (2018).

63. D. Wang, J. Wengrod, L. B. Gardner, Overexpression of the c-myc oncogene inhibits nonsense-mediated RNA decay in B lymphocytes. J Biol Chem 286, 40038–40043 (2011).

64. J. Wengrod et al., Inhibition of nonsense-mediated RNA decay activates autophagy. Mol Cell Biol 33, 2128–2135 (2013).

65. J. Ye et al., The GCN2-ATF4 pathway is critical for tumour cell survival and proliferation in response to nutrient deprivation. EMBO J 29, 2082–2096 (2010).

66. C. J. Liu et al., OEPR Cloning: an Efficient and Seamless Cloning Strategy for Large- and Multi-Fragments. Sci Rep 7, 44648 (2017).

67. A. Dobin et al., STAR: ultrafast universal RNA-seq aligner. Bioinformatics 29, 15–21 (2013).

68. B. Li, C. N. Dewey, RSEM: accurate transcript quantification from RNA-Seq data with or without a reference genome. BMC Bioinformatics 12, 323 (2011).

69. K. Wang, M. Li, H. Hakonarson, ANNOVAR: functional annotation of genetic variants from high-throughput sequencing data. Nucleic Acids Res 38, e164 (2010).

70. M. I. Love, W. Huber, S. Anders, Moderated estimation of fold change and dispersion for RNA-seq data with DESeq2. Genome Biol 15, 550 (2014).

71. S. E. Harvey, C. Cheng, Methods for Characterization of Alternative RNA Splicing. Methods Mol Biol 1402, 229–241 (2016).

72. S. M. Blue et al., Transcriptome-wide identification of RNA-binding protein binding sites using seCLIP-seq. Nat Protoc 17, 1223–1265 (2022).

73. S. Heinz et al., Simple combinations of lineage-determining transcription factors prime cis-regulatory elements required for macrophage and B cell identities. Mol Cell 38, 576–589 (2010).

74. D. Y. Wu, D. Bittencourt, M. R. Stallcup, K. D. Siegmund, Identifying differential transcription factor binding in ChIP-seq. Front Genet 6, 169 (2015).

